# Piezo1-dependent regulation of pericyte proliferation by blood flow during brain vascular development

**DOI:** 10.1101/2023.05.01.538849

**Authors:** Huaxing Zi, Xiaolan Peng, Jianbin Cao, Jiwen Bu, Jiulin Du, Jia Li

## Abstract

Blood flow is known to regulate cerebrovascular development through acting on vascular endothelial cells (ECs). As an indispensable component of the cerebrovascular unit, brain pericytes physically couple with ECs with the highest density in the body and play vital roles in blood-brain barrier integrity maintenance and neurovascular coupling. However, it remains unclear whether blood flow affects the development of brain pericytes. Here, we report that blood flow can promote brain pericyte proliferation, which depends on the mechanosensitive ion channel Piezo1. Using *in vivo* time-lapse imaging of larval zebrafish, we monitored the developmental dynamics of brain pericytes and found that they proliferated to expand their population and increase their coverage on brain vessels. In combination with pharmacological and genetic approaches, we demonstrate that the proliferation of brain pericytes can be enhanced by increased blood flow through Piezo1 expressed in ECs. Furthermore, EC-intrinsic Notch signaling was found to be downstream of Piezo1 for the blood flow regulation of brain pericyte proliferation. Thus, our findings reveal an important role of blood flow for pericyte proliferation, extending the functional spectrum of hemodynamics on the development of cerebral vasculature.

## Introduction

Pericytes, the mural cells of small vessels, tightly associate with ECs by forming multiple types of intercellular junctions, allowing them act as a physical integration in response to mechanical and chemical stimuli ***(Armulik et al., 2011***). In the brain, pericytes cover capillaries with the highest density in the body and are necessary for diverse brain-specific vascular functions (***Winkler et al., 2011***). To maintain the hemostasis of the brain and support neural activities, brain vessels are specialized to form blood-brain barrier (BBB), which restricts the permeability of circulating substrates (***Zhao et al., 2015***). As a cellular component of BBB, brain pericytes are important for BBB integrity during development (***Armulik et al., 2010; Daneman et al., 2010***). Brain pericytes also participate in neurovascular coupling (NVC), by which local blood flow speeds up in response to neural activities (***Alarcon-Martinez et al., 2020; Kisler et al., 2017; Mishra et al., 2016***). Moreover, pericyte loss was found to be associated with neural disorders, including Alzheimer’s disease and vascular dementia (***Liu et al., 2010; Ross Nortley, 2019; Sagare et al., 2013; Yang et al., 2022; Yao et al., 2023***). Therefore, it is of importance to understand the mechanism governing the population expansion and maintenance in the brain during development and adult stage.

During development, brain pericyte progenitors emerge from pericerebral vessels, and then migrate into the brain along vessels, a process depended on PDGFB-PDGFRb signaling (***Ando et al., 2016; Lindahl et al., 1997***). After ingress into the brain, pericytes can proliferate, which is regulated cell-autonomously by Notch signaling and possibly PDGFRb signaling in pericytes (***Ando et al., 2016; Wang et al., 2014***). Additionally, the high density and specialized functions of pericytes in the brain suggest that brain environment, such as the distinct hemodynamics of cerebral vasculature (***Chen et al., 2012***), may play a role in brain pericyte development.

Responding to physiological mechanics is an intrinsic property of cells, which is critical for their normal functioning. Blood flow constantly generates mechanical forces on vascular endothelial wall and plays crucial roles for vascular development (***Hahn and Schwartz, 2009; le Noble et al., 2008***). Blood flow is known to regulate the arteriovenous differentiation via activating higher Notch signaling in committed arterial ECs (***Geudens et al., 2019; Lasch et al., 2019; le Noble et al., 2004; Obi et al., 2009***). During cerebrovascular patterning, blood flow promotes vessel maintenance via the nuclear import of YAP (***Nakajima et al., 2017***), and decrease of blood flow drives the pruning of blood vessels (***Chen et al., 2012***). Moreover, the high pericyte coverage is a unique feature of brain vasculature, which was thought potentially due to the effect of the distinct hemodynamics of the cerebral vasculature (***Dessalles et al., 2021***), but the underlying mechanism has not been probed. Considering the functions of brain pericytes in blood flow regulation and the tight physical association between pericytes and ECs (***Armulik et al., 2011***), it is crucial to investigate whether blood flow could regulate the development of pericytes in the brain.

In this study, we thus investigated the role of blood flow in brain pericyte development by using the zebrafish model, and found that blood flow can promote brain pericyte proliferation during vascular development. Through *in vivo* time-lapse imaging, we observed that blood flow positively regulates the population and coverage of brain pericytes by promoting their proliferation. By combining genetic and pharmacological manipulations, we proved that this effect depends on the activation of the mechanosensitive ion channel Piezo1 in ECs. Further investigation revealed found that the Piezo1 activation could up-regulate Notch signaling in ECs. Enhancing EC-specific Notch activation promoted pericyte proliferation. On the contrary, EC-specific Notch inhibition suppressed Piezo1 activation-induced pericyte proliferation. These results indicate that Notch signal acts as the downstream effector of Piezo1 in ECs for brain pericyte proliferation. Thus, our study identifies a cascade of blood flow-Piezo1-Notch signaling in ECs important for brain pericyte proliferation, revealing a novel role of blood flow in regulating the development of perivascular cells in the brain.

## Results

### Pericytes expand their population by proliferation after ingress into the brain

To *in vivo* monitor pericyte dynamics during development in zebrafish, we targeted the gene of zebrafish Pdgfrb, a commonly used marker for pericytes (***Ando et al., 2016; Lindahl et al., 1997***), and generated a *Ki(pdgfrb:GAL4-VP16)* line by utilizing CRISPR/Cas9-mediated knock-in (***Li et al., 2015***). The *Ki(pdgfrb:GAL4-VP16)* line expresses Gal4-VP16 under the control of the endogenous *pdgfrb* promoter (***Figure 1-figure supplementary 1A, B***). To visualize the brain pericytes, we performed *in vivo* imaging of *Ki(pdgfrb:GAL4-VP16);Tg(4×nrUAS:GFP);Tg(fli1a:DsRedEx)* larvae, in which pericytes were labeled with green fluorescent protein (GFP) and ECs were labeled with the red fluorescent protein DsRed (***Figure 1-figure supplementary 1C***). And we observed that the GFP-expressing cells wrapped the blood vessels with their long fibers in the brain (***Figure 1A***), which is the typical morphology of pericytes (***Ando et al., 2016***), indicating the efficient labeling of pericytes in this line.

**Figure 1.**
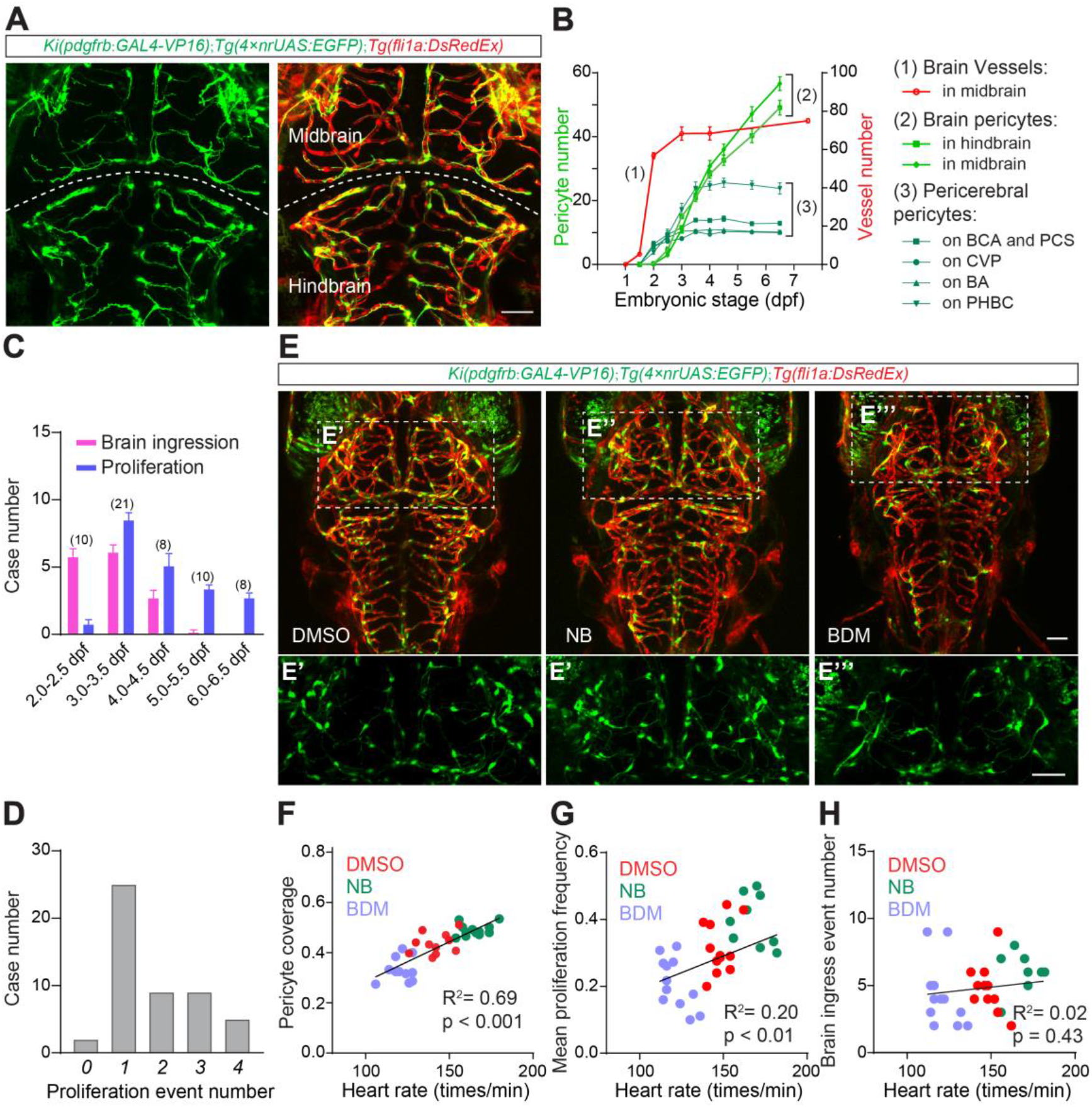
Blood flow promotes brain pericyte proliferation. (A) Representative images of the pericytes and blood vessels in the brain of the *Ki(pdgfrb:GAL4-VP16);Tg(4×nrUAS:GFP);Tg(fli1a:DsRedEx)* larvae at 4.5 dpf. The images are partially z-projected. The merged and single channel images with endothelial cells labeled in red and pericytes labeled in green are shown here and hereafter. (B) Graph showing the numerical changes of the blood vessels in the midbrain, the pericytes in the brain and the pericytes on the surrounding vessels (BCA&PCS, CVP, BA and PHBC). N=9 for pericytes, and n=8 for midbrain vessels. (C) Graphic showing the event numbers of the ingress of pericerebral pericytes into the brain and the proliferation of brain pericytes over a 12-hour period each day in the early embryonic stage. (D) Graphic showing the number of cases in which a photoconverted single brain pericyte proliferated for varying times between 2.5 and 4.5 dpf. (E) Representative images of the pericytes and blood vessels in the brain of the DMSO, NB and BDM treated larvae at 4.5 dpf. The images of pericytes in the midbrain are highlighted in (E’)-(E’’’). (F-H) Linear regression analysis for effects of pharmacological blood flow manipulations on pericyte coverage of brain vessels, brain pericyte proliferation and the ingress of the pericerebral pericytes into the brain. DMSO, NB and BDM are applied during 3.5-4.5 dpf for (F) and (G), and 3.0-3.5 dpf for (H) and (I). Images are shown from a top view. Scale bar, 50 μm for (A) and (E). Images of 4.5 dpf larvae are used for counting the number of brain pericytes and the pericyte coverage of brain vessels, time-lapse images during 3.0-3.5 dpf are used for counting the event numbers of brain pericyte proliferation and ingress of the pericerebral pericytes into the brain. Data are represented as mean ± SEM. N values are shown above the bars in (C). Stars represent the results of unpaired two-tailed Student’s t-test between groups (*p < 0.05, **p < 0.01, ***p < 0.001, ****p < 0.0001). See also ***Figure 1—figure supplementary 1-3***.

To study the dynamics of brain pericytes during cerebrovascular development, we carried out long-term serial confocal imaging of *Ki(pdgfrb:GAL4-VP16);Tg(4×nrUAS:GFP);Tg(fli1a:DsRedEx)* larvae during 1.5 - 6.5 days post-fertilization (dpf) (***Figure 1-figure supplementary 2A***). We found that, at 1.5 dpf, no pericyte was present in the brain or on the pericerebral vessels, although the cerebral vessels had already begun forming (***Figure 1B*** and ***Figure 1-figure supplementary 2A***). Pericytes started to appear on pericerebral vessels at around 2.0 dpf and were initially observed in the brain at around 2.5 dpf (***Figure 1B*** and ***Figure 1-figure supplementary 2A***). After 2.5 dpf, the number of pericytes markedly increased in the midbrain and hindbrain, and the brain vasculature was fully covered by pericytes at 6.5 dpf (***Figure 1B*** and ***Figure 1-figure supplementary 2A***).

To determine how the number of pericytes increase in the brain, we performed *in vivo* time-lapse imaging with the interval of 2 - 4 hours during 2.0 - 6.5 dpf. We found that pericytes on pericerebral vessels, including the basal communicating artery (BCA), choroidal vascular plexus (CVP), posterior communicating segment (PCS) and basilar artery (BA), migrated into the brain along perfused blood vessels (***Figure 1C*** and ***Figure 1-figure supplementary 2B***). After ingress to the brain, pericytes underwent proliferation to expand their population (***Figure 1C*** and ***Figure 1-figure supplementary 2C***). The ingression of pericerebral pericytes was the major contribution to the growth of brain pericyte population during 2.0 - 2.5 dpf (***Figure 1C***). While the proliferation of pericytes in the brain took a dominant role after 3.0 - 3.5 dpf and the ingress of pericerebral pericytes could hardly be observed after 5.0 dpf (***Figure 1C***). Therefore, brain pericytes expand their population mainly through proliferation after ingress to the brain during development.

To further investigate the proliferation capability of brain pericytes, we tracked the fate of specific brain pericytes during embryogenesis by using the *Ki(pdgfrb:GAL4-VP16);Tg(UAS:Kaede)* line, which expresses the stable photoconvertible Kaede protein in pericytes ***(Schuster and Ghysen, 2013***). We photoconverted a single brain pericyte in each larva from green to red at 2.5 dpf, and then examined the number of pericytes with red fluorescence to calculate the number of proliferation events in the brain at 4.5 dpf (***Figure 1D*** and ***Figure 1-figure supplementary 2D***). We found that 48 out of 50 photoconverted brain pericytes underwent cell division during 2.5 – 4.5 dpf (***Figure 1D***). Moreover, 22 out of 50 photoconverted brain pericytes divided more than once (***Figure 1D***). To further confirm the presence of pericyte proliferation in the brain, we visualized the cycling brain pericytes using the *Tg(EF1a1:mAG-gmnn.100);Ki(pdgfrb:GAL4-VP16);Tg(UAS:mCherry-MA)* line, which allows for detection of the cells in S/G2/M phase by green fluorescence based on the fluorescent ubiquitination-based cell cycle indicator (FUCCI) system (***Sugiyama et al., 2009***). We observed that about 30% of the pericytes in brain displayed green fluorescence at 3.5 dpf (***Figure 1-figure supplementary 2E***), indicating the proliferating state of these cells. These results demonstrate that pericytes frequently proliferate, leading to the constant expansion of their population in the brain during the early embryonic stage.

### Blood flow promotes the proliferation of brain pericytes

Having characterized the developmental signatures of the brain pericytes during embryogenesis, we next investigated the role of blood flow for brain pericyte development. We used the heart rate as a proxy for blood flow velocity (***Chen et al., 2012***), and evaluated the population density of brain pericytes by calculating the pericyte coverage on brain vessels (***Xu et al., 2017***). To change the velocity of blood flow, the *Ki(pdgfrb:GAL4-VP16);Tg(4×nrUAS:GFP);Tg(fli1a:DsRedEx)* larvae were treated with norepinephrine bitartrate (NB) or 2,3-butanedione-2-monoxime (BDM) to up-regulate or down-regulate their heart rates during 3.5 – 4.5 dpf, respectively (NB = 164.7±2.2 versus DMSO = 141.8±9.6, increased by 16%, p < 0.0001; BDM = 121.1±6.9 versus DMSO = 141.8±9.6, decreased by 15%, p < 0.0001) (***Chen et al., 2012***) (***Figure 1-figure supplementary 3A***). We performed *in vivo* imaging and recorded the heart rate of the larvae simultaneously at 4.5 dpf (***Figure 1E***), and found that the pericyte coverage of brain vessels was positively correlated with the heart rate (liner-regression: R^2^ = 0.69, p<0.001) (***Figure 1F***). NB treated larvae displayed significantly increased pericyte coverage on brain vessels (NB = 0.49±0.01 versus DMSO = 0.43±0.01, increased by 14%, p < 0.001), and BDM application caused significant reduction of the pericyte coverage (BDM = 0.33±0.01 versus DMSO = 0.43±0.01, decreased by 23%, p < 0.0001) (***Figure 1-figure supplementary 3B***). These results indicate that blood flow plays a promotional role in the population expansion of brain pericytes during embryogenesis.

Next, we then investigated how blood flow increased the pericyte coverage on brain vessels. Previous data showed that both of the pericyte ingress into the brain and the brain pericyte proliferation could contribute to the pericyte number increase in the brain. We therefore performed long-term time-lapse imaging to discriminate which of the ways could be regulated by blood flow during cerebrovascular development. We found that, during 3.0 - 3.5 dpf, the mean proliferation frequency of brain pericytes positively relate to the heart rate (liner-regression: R^2^ = 0.2, p<0.01) ***(Figure 1G***). The NB treated larvae displayed significant increase of brain pericyte proliferation frequency (NB = 0.39±0.02 versus DMSO = 0.32±0.02, increased by 22%, p < 0.05), and BDM markedly decreased the brain pericyte proliferation frequency (BDM = 0.21±0.02 versus DMSO = 0.32±0.02, decreased by 34%, p < 0.01) (***Figure 1-figure supplementary 3C***). Whereas the pericyte ingress into the brain was not significantly affected (R^2^ = 0.02, p = 0.43 for liner-regression analysis; NB versus DMSO, p = 0.57; BDM versus DMSO, p = 0.57) (***Figure 1H*** and ***Figure 1-figure supplementary 3D***). These findings indicate that blood flow positively regulates the population expansion of brain pericytes by enhancing their proliferation.

### EC-specific Piezo1 mediates the effect of blood flow on brain pericyte proliferation

As ECs line between the blood and pericytes, and they can directly sense the dynamics of blood flow, we therefore sought to identify the mechanical sensor expressed in ECs that is responsible for blood flow regulation of brain pericyte proliferation. Piezo1, a mechanosensitive ion channel expressed on the membrane of ECs, plays crucial roles in EC alignment and angiogenesis during vascular development (***Li et al., 2014; Liu et al., 2020; Ranade et al., 2014***). Furthermore, Piezo1 was recently identified as the mechanosensor in brain vessels, and loss-of-function of which blocked the mechanical force-triggered Ca^2+^ signals in ECs (***Harraz et al., 2022***). Therefore, we asked whether Piezo1 plays a role for brain pericyte proliferation.

We used the *piezo1* knockout line *piezo1^ion89d^* (abbreviated as *piezo1^-/-^*) to investigate the involvement of Piezo1 in brain pericyte proliferation ***(Liu et al., 2020***). By performing *in vivo* imaging at 4.5 dpf, we found that the *piezo1^-/-^* larvae displayed significantly reduced brain pericyte number (*piezo1^-/-^* = 57±3.6 versus WT = 71±1.5, decreased by 20%, p < 0.001) and pericyte coverage of brain vessels (*piezo1^-/-^ =* 0.34±0.02 versus WT = 0.45±0.01, decreased by 24%, p < 0.0001) compared to the sibling wild type (WT) larvae (***Figure 2A, B*** and ***Figure 2-figure supplementary 1A***), demonstrating the requirement of Piezo1 for the expansion of brain pericyte population. Moreover, by performing long-term time-lapse imaging during 3.0 - 3.5 dpf, we found that the *piezo1* knockout larvae displayed reduced proliferation frequency of brain pericytes (*piezo1^-/-^* = 0.19±0.03 versus WT = 0.33±0.03, decreased by 42%, p < 0.01) (***Figure 2F***). These results demonstrate that Piezo1 is required for brain pericyte proliferation.

**Figure 2.**
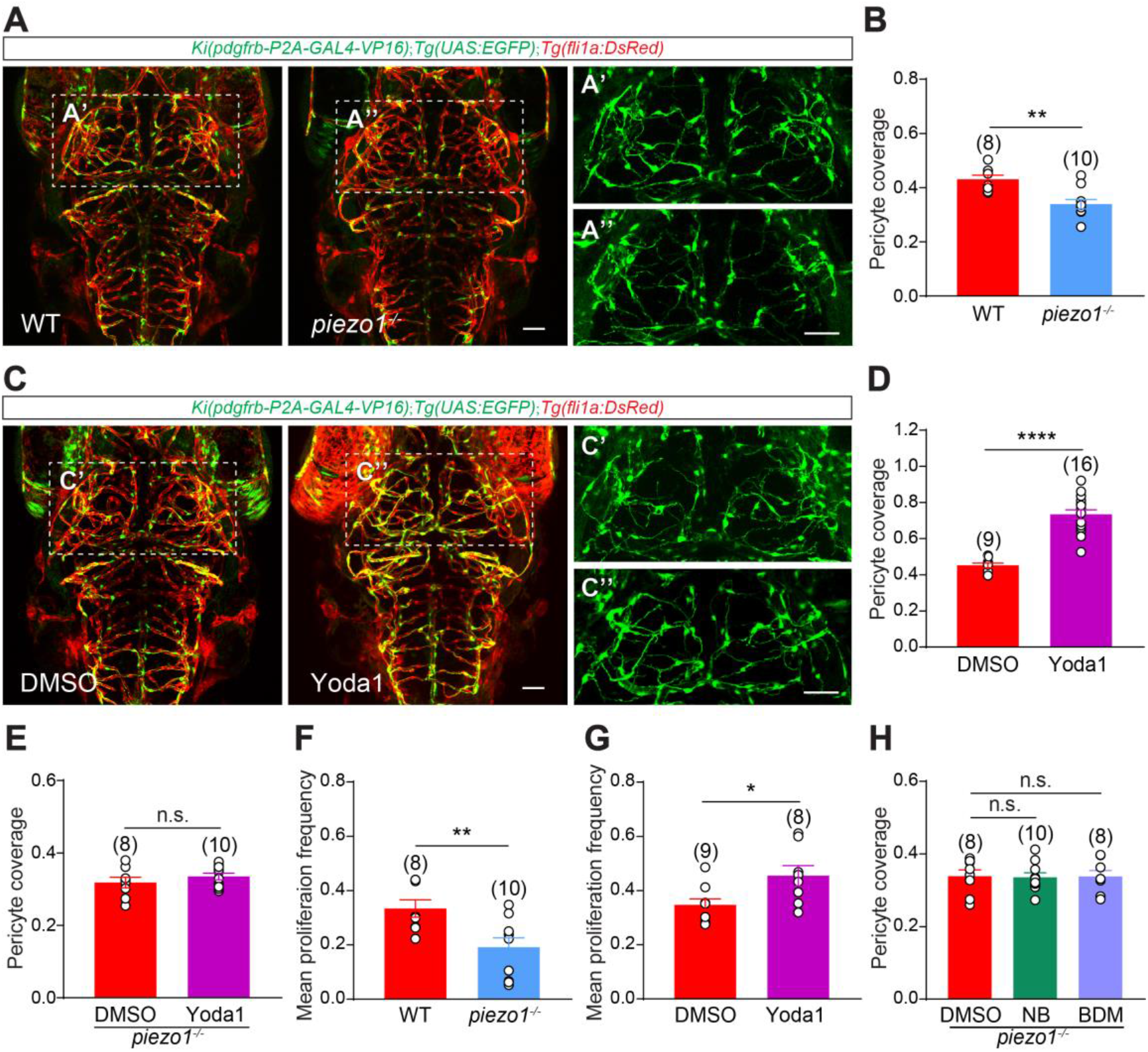
Piezo1 mediates the effect of blood flow on brain pericyte proliferation. (A) Representative images of pericytes and blood vessels in the midbrain of wild type (WT) and Piezo1-knockout (*piezo1^-/-^*) larvae at 4.5 dpf. The images of pericytes in the midbrain are highlighted in (A’) and (A’’). (B) Summary of the effect of *piezo1* knockout on pericyte coverage of brain vessels. (C) Representative images of pericytes and blood vessels in the brain of control (DMSO) and Piezo1 activator (Yoda1)-treated larvae at 4.5 dpf. The images of pericytes in the midbrain are highlighted in (C’) and (C’’). (D) Summary of the effect of Yoda1 treatment on pericyte coverage of brain vessels. (E) Summary of the effect of Yoda1 on the pericyte coverage of brain vessels in *piezo1^-/-^* larvae. (F) Summary of the effect of *piezo1* knockout on brain pericyte proliferation. (G) Summary of the effect of Yoda1 treatment on brain pericyte proliferation. (H) Summary of the effects of NB and BDM treatment on pericyte coverage of brain vessels in *piezo1^-/-^*larvae. Images are shown from a top view. Scale bar, 50 μm for (A) and (C). Images of 4.5 dpf larvae are used for counting the brain pericyte number and the pericyte coverage of brain vessels, time-lapse images during 3.0-3.5 dpf are used for counting the event numbers of brain pericyte proliferation. Data are represented as mean ± SEM. N values are shown above the aligned plots. Stars represent the results of unpaired two-tailed Student’s t-test between groups (*p < 0.05, **p < 0.01, ***p < 0.001, ****p < 0.0001). See also ***Figure 2—figure supplementary 1***.

Next, we asked: whether increasing Piezo1 activity could promote brain pericyte proliferation? We treated the larvae with Yoda1, an agonist of Piezo1 (***Botello-Smith et al., 2019***), and observed suppressed cerebrovascular pattern in the Yoda-treated larvae (***Figure 2C***), consistent with our previously reported function of Piezo1 in cerebrovascular development (***Liu et al., 2020***). Although the number of brain pericytes in the Yoda1-treated larvae did not significantly up-regulated compared to the DMSO treated controls, the pericyte coverage of brain vessels in these larvae significantly increased (Yoda1 = 0.73±0.03 versus DMSO = 0.43±0.01, increased by 70%, p < 0.0001) (***Figure 2C, D***). In contrast, application of Yoda1 had no effect on the pericyte coverage of brain vessels in the *piezo1^-/-^* larvae (Yoda1 versus DMSO, p = 0.84) (***Figure 2E***), confirming that Yoda1-induced effect on brain pericytes was dependent on Piezo1. Consistently, by using time-lapse imaging during 3.0 - 3.5 dpf, we found that the brain pericytes in Yoda1-treated larvae displayed significantly increased proliferation frequency (Yoda1 = 0.45±0.04 versus DMSO = 0.35±0.02, increased by 29%, p < 0.05) (***Figure 2G***). In addition, neither knockout of piezo1 nor application of Yoda1 could significantly change the heart rate of the larvae (***Figure 2-figure supplementary 1C***). Thus, increasing Piezo1 activity can promote brain pericyte proliferation.

As Piezo1 is expressed in multiple types of cells in the brain (***Vanlandewijck et al., 2018; Yang et al., 2022***), we sought to investigate the role of EC-specific Piezo1 in brain pericyte development. To achieve this, we generated a *Ki(piezo1-loxp)* zebrafish line, which allowed us to specifically knock out *piezo1* in the ECs by crossing to a EC-specific Cre-expressing line *Tg(flk1:Cre)* (abbreviated as *piezo1^ΔEC^*) (***Figure 3-figure supplementary 1A-C***) (***Li et al., 2015; Pan et al., 2005***). We confirmed the efficient deletion of the exon 24 of *piezo1* in the *piezo1^ΔEC^*larvae by PCR and sequencing (***Figure 3-figure supplementary 1D, E***).

To examine the effect of EC-specific *piezo1* knockout on brain pericyte development, we performed *in vivo* imaging of the *piezo1^ΔEC^* and the sibling wild-type larvae (***Figure 3A***), and found that the *piezo1^ΔEC^*larvae displayed significant reduced brain pericyte number and pericyte coverage of brain vessels (number: *piezo1^ΔEC^* = 58.4±2.4 versus WT = 71.0±2.8, decreased by 18%, p < 0.01; coverage: *piezo1^ΔEC^* = 0.33±0.01 versus WT = 0.45±0.01, decreased by 18%, p < 0.001) (***Figure 3A-C***). Meanwhile, the brain pericyte number and the pericyte coverage of brain vessels in the *Ki(piezo1-loxp)^+/+^* (abbreviated as *piezo1^f/f^*) larvae were similar to the sibling wild-type controls (number: *piezo1^f/f^* versus WT, p = 0.59; coverage: *piezo1^f/f^* versus WT, p = 0.70) (***Figure 3A-C***). Moreover, by using long-term *in vivo* time-lapse imaging, we found that the *piezo1^ΔEC^* larvae exhibited significantly decreased brain pericyte proliferation frequency (*piezo1^ΔEC^* = 0.16±0.03 versus *piezo1^f/f^* = 0.32±0.03, decreased by 50%, p < 0.01), in comparison with the sibling *piezo1^f/f^* larvae (***Figure 3D***). These results indicate that EC-specific Piezo1 is required for brain pericyte proliferation during vascular development.

**Figure 3.**
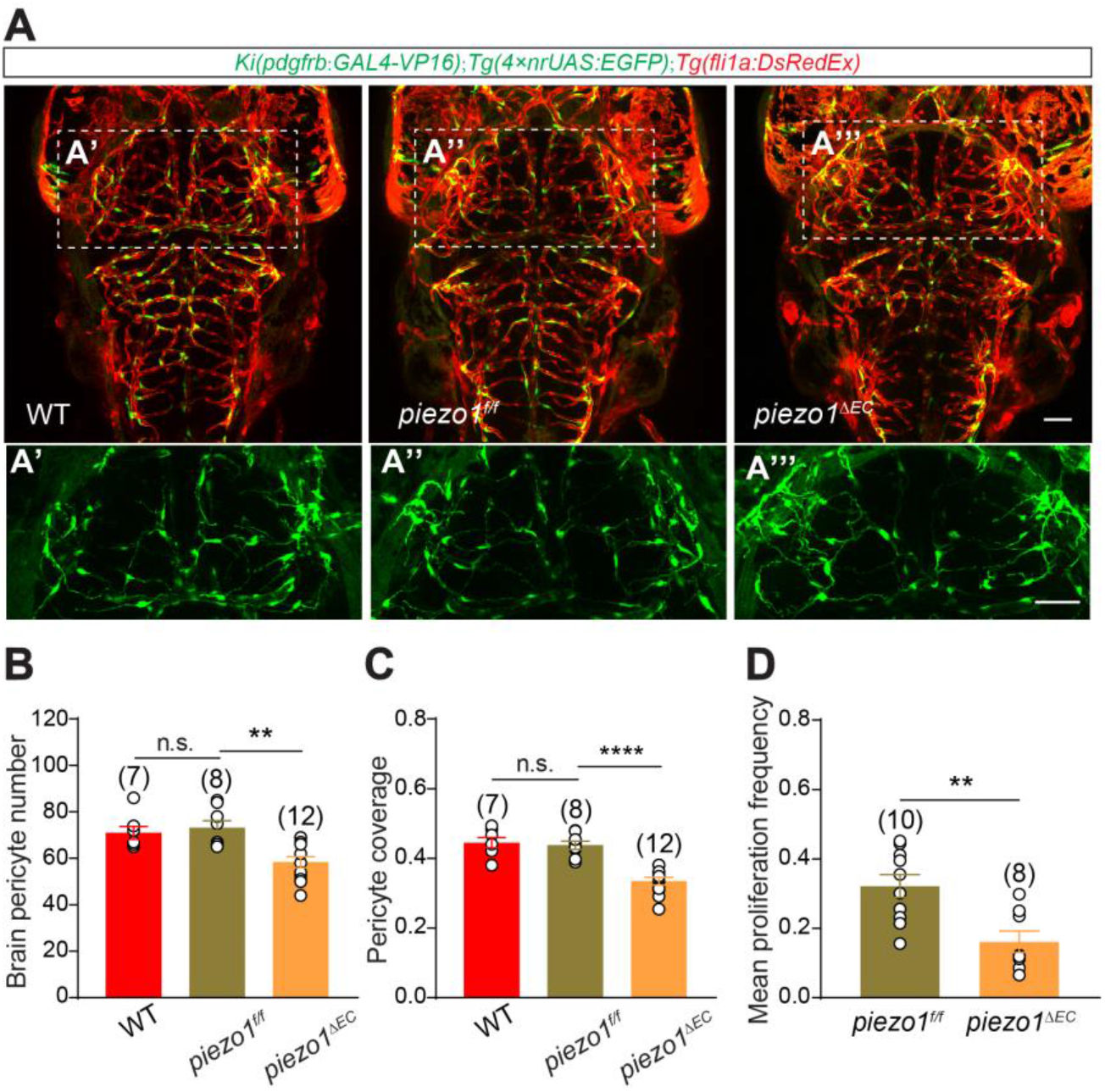
EC-specific Piezo1 is required for brain pericyte proliferation. (A) Representative images of the pericytes and blood vessels in the midbrain of WT and *piezo1^f/f^*and *piezo1^ΔEC^* larvae at 4.5 dpf. The images of pericytes in the midbrain are highlighted in (A’)-(A’’’). (B-D) Summary of the effect of EC-specific Piezo1 knockout on brain pericyte number (B), pericyte coverage of brain vessels (C), and brain pericyte proliferation (D), compared to the sibling *piezo1^f/f^* larvae used as controls. Images are shown from a top view. Scale bar, 50 μm for (A) and (A’)-(A’’’). Images of 4.5 dpf larvae are used for counting the brain pericyte number and the pericyte coverage of brain vessels, time-lapse images during 3.0-3.5 dpf are used for counting the event number of brain pericyte proliferation. Data are represented as mean ± SEM. N values are shown above the aligned plots. Stars represent the results of unpaired two-tailed Student’s t-test between groups (*p < 0.05, **p < 0.01, ***p < 0.001, ****p < 0.0001). See also ***Figure 3—figure supplementary 1***.

The *piezo1^ΔEC^* larvae displayed the similar phenotypes of pericytes and vessels in the brain as the *piezo1^-/-^* larvae, suggesting that Piezo1 regulates brain pericyte proliferation in a tissue-specific manner. To probe whether blood flow relies on Piezo1 to regulate brain pericyte proliferation, we changed the blood flow velocity on the *piezo1* knockout larvae by NB or BDM (***Figure 2-figure supplementary 1D***), and found that neither of them could change the pericyte coverage of brain vessels (NB versus DMSO, p = 0.87; BDM versus DMSO, p = 0.97) (***Figure 2H*** and ***Figure 1-figure supplementary 3B***)). Therefore, these results indicate that Piezo1 mediates the effect of blood flow on brain pericyte proliferation.

### Notch signaling functions as the downstream of Piezo1 in ECs for brain pericyte proliferation

Next, we asked: what is the downstream target of Piezo1 in ECs that is required for brain pericyte proliferation? It is reported that Notch signaling in ECs could be up-regulated by the blood flow in a Piezo1-dependent manner *in vitro* and *in vivo* (***Caolo et al., 2020; Duchemin et al., 2019***). To test whether Piezo1 can change Notch signaling *in vivo*, we increased Piezo1 activity on the *Tg(Tp1:d2GFP*) Notch reporter line, which expresses a destabilized green fluorescent protein upon Notch activation (***Clark et al., 2012***) (***Figure 4-figure supplementary 1A***). We found that Yoda1 treatment markedly increased the number of d2GFP-expressing ECs in the brain (Yoda1 = 4.8±0.65 versus DMSO = 0.6±0.27, increased by 700%, p < 0.001) (***Figure 4A, B***). And the increase could be suppressed by the Piezo1 inhibitor GsMTx4 (Yoda1+GsMTx4 = 1.46±0.28 versus Yoda1 = 4.80±0.65, decreased by 70%, p < 0.01) (***Liu et al., 2020***) (***Figure 4B***), suggesting that Piezo1 could regulate Notch activity in brain ECs during development.

**Figure 4.**
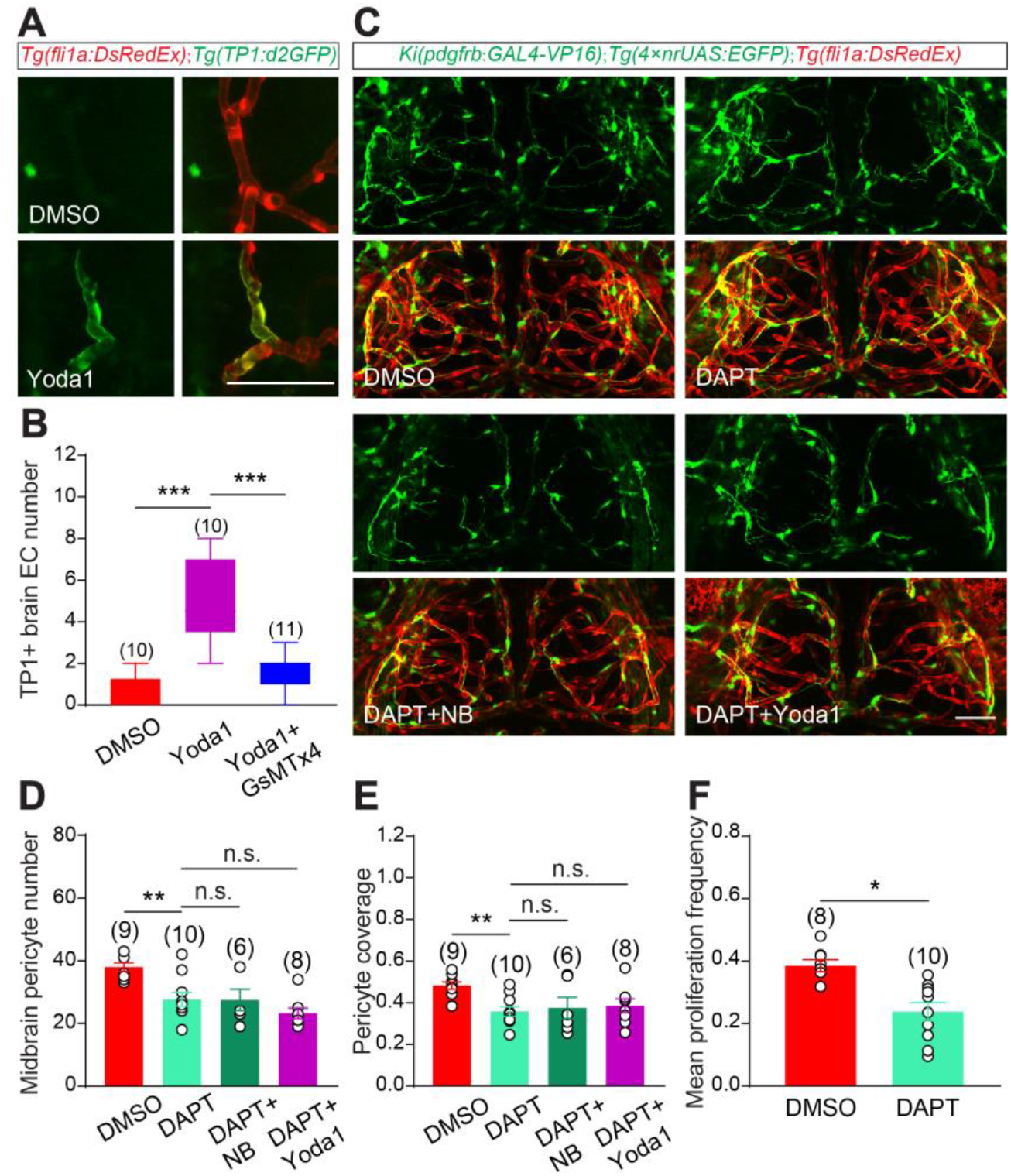
Notch signaling functions as the downstream of Piezo1 in regulating pericyte proliferation. (A) Representative images of the d2GFP-labeled brain ECs with Notch activation in theYoda1-treated *Tg(fli1a:DsRedEx);Tg(Tp1:d2GFP)* larvae at 4.5 dpf, compared to the location-matched ECs in the DMSO control larvae. (B) Summary of the effect of Yoda1 and GsMTx4 on the Notch signaling in brain ECs. (C) Representative images of the pericytes and blood vessels in the midbrain of DMSO, DAPT, DAPT+NB and DAPT+Yoda1 treated larvae at 4.5 dpf. (D) Summary of the effects of DAPT, DAPT+NB and DAPT+Yoda1 treatment on the number of pericytes in the midbrain. (E) Summary of the effects of DAPT, DAPT+NB and DAPT+Yoda1 treatment on pericyte coverage of vessels in the midbrain. (F) Summary of the effect of DAPT treatment on the brain pericyte proliferation. Images are shown from a top view. Scale bar, 50 μm for (A) and (C). Images of 4.5 dpf larvae are used for counting the number of pericytes and the pericyte coverage of the vessels in the midbrain, time-lapse images during 3.0-3.5 dpf are used for counting the event numbers of brain pericyte proliferation. Data are represented as mean ± SEM. The n values are shown above the aligned plots. Stars represent the results of unpaired two-tailed Student’s t-test between groups (*p < 0.05, **p < 0.01, ***p < 0.001, ****p < 0.0001). See also ***Figure 4—figure supplementary 1***.

Notch3 is highly expressed in pericyte and is critically important for pericyte development (***Wang et al., 2014***). However, whether EC-intrinsic Notch signaling could regulate brain pericyte proliferation and mediate the effect of blood flow on brain pericyte development is still unclear. Therefore, to determine the involvement of Notch signaling in blood flow regulation of brain pericyte proliferation, we first used the γ-secretase inhibitor DAPT to inhibit Notch signaling on the blood flow or the Piezo1 activity increased embryos (***Geling et al., 2002***) (***Figure 4C***). We found that Notch inhibition significantly decreased the number of brain pericytes, the pericyte coverage of brain vessels and the brain pericyte proliferation frequency (number: DAPT = 27.7±2.21 versus DMSO = 35.4±1.18, decreased by 22%, p < 0.01; coverage: DAPT = 0.36±0.02 versus DMSO = 0.43±0.01, decreased by 16%, p < 0.01; proliferation frequency: DAPT = 0.24±0.03 versus DMSO = 0.31±0.02, decreased by 23%, p < 0.05) (***Figure 4C-F***). However, neither NB nor Yoda1 could change the number of brain pericytes or pericyte coverage of brain vessels in the present of DAPT (number: DAPT+NB = 27.5±3.4 versus DAPT = 38±1.3, p = 0.96; coverage: DAPT+NB =0.37±0.05 versus DAPT = 0.36±0.02, p = 0.49) (***Figure 4C-E***), indicating that blood flow-Piezo1 relies on Notch signaling to regulate brain pericyte proliferation.

To examine the role of EC-intrinsic Notch signaling in brain pericyte proliferation, we next genetically manipulated the Notch signaling in ECs by generating T*g(flk1:H2B-mNeoGreen-dnMAML)* and *Tg(flk1:H2B-mScarlet-NICD)* fish lines to decrease or increase Notch signaling in ECs, respectively (***Galvez-Santisteban et al., 2019; Samsa et al., 2015***). In comparison with the controls, the *Tg(flk1:H2B-mNeoGreen-dnMAML)* larvae, in which ECs express the green fluorescent protein mNeoGreen and the dominant negative Mastermind-like (dnMAML) (***Figure 5-figure supplementary 1A***), displayed significantly decreased number of pericytes and pericyte coverage in the midbrain (number: dnMAML = 24.5±2.3 versus control = 34.0±1.1, decrease 28%, p < 0.01; coverage: dnMAML = 0.31±0.10 versus control = 0.46±0.07, decreased by 33%, p < 0.001) (***Figure 5A-C***). While, the *Tg(flk1:H2B-mScarlet-NICD)* larvae, in which ECs express the intercellular domain of Notch1 (NICD) and the red fluorescent protein mScarlet (***Figure 5-figure supplementary 1B***), showed increased number of brain pericytes and pericyte coverage of brain vessels (number: NICD = 46.4±4.0 versus control = 34.0±1.1, increased by 36%, p < 0.05; coverage: NICD versus control: 1.0±0.34 versus 0.46±0.07, increased by 117%, p < 0.0001) (***Figure 5A-C***). Furthermore, long-term time-lapse imaging showed that expressing dnMAML in ECs significantly reduced the proliferation frequency of brain pericytes (dnMAML = 0.24±0.03 versus control = 0.32±0.09, decreased by 25%, p < 0.001), while expressing NICD in ECs markedly increased the brain pericyte proliferation frequency (NICD = 0.50±0.09 versus control = 0.31±0.09, increased by 61%, p < 0.001) (***Figure 5D***). These findings indicate that EC-intrinsic Notch signaling can positively regulate the population expansion of brain pericytes by promoting their proliferation.

**Figure 5.**
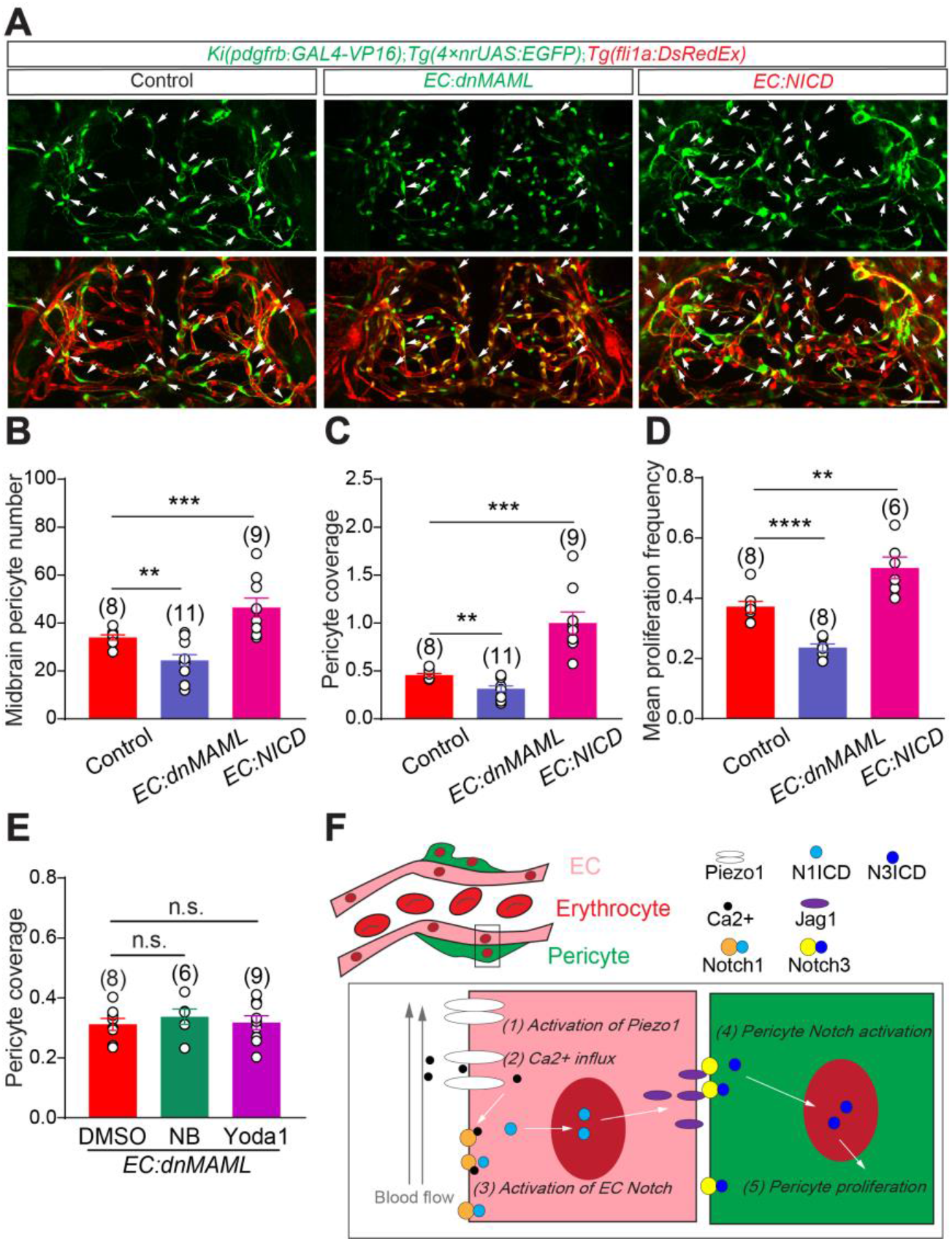
EC-specific Notch signaling mediates blood flow regulation of brain pericyte proliferation. (A) Representative images of the pericytes and blood vessels in the midbrain of the control, *Tg(flk1:tetoff-H2B-mNeoGreen-P2A-dnMAML* (abbreviated to *EC: dnMAML*), and *Tg(flk1: tetoff-H2B-mScarlet-P2A-NICD)*(abbreviated to *EC:NICD*) larvae at 4.5 dpf. (B) Summary of the effects of EC specific Notch activation and inhibition on the number of pericytes in the midbrain. (C) Summary of the effects of EC specific Notch signaling activation and inhibition on pericyte coverage of vessels in the midbrain. (D) Summary of the effects of EC specific Notch activation and inhibition on brain pericyte proliferation. (E) Summary of the effects of DMSO, NB and Yoda1 treatment on pericyte coverage of vessels in the midbrain of the *Tg(flk1:tetoff-H2B-mNeoGreen-P2A-dnMAML)* larvae. (F) Our working model. Images are shown from a top view. Scale bar, 50 μm for (A). Images of 4.5 dpf larvae are used for counting the number of pericytes and the pericyte coverage of the vessels in the midbrain, time-lapse images during 3.0-3.5 dpf are used for counting the event numbers of brain pericyte proliferation. Data are represented as mean ± SEM. The n values are shown above the aligned plots. Stars represent the results of unpaired two-tailed Student’s t-test between groups (*p < 0.05, **p < 0.01, ***p < 0.001, ****p < 0.0001).See also ***Figure 5—figure supplementary 1***.

To determine whether blood flow regulates brain pericyte proliferation specifically through Notch signaling in ECs, we increased blood-flow or piezo1 activity to determine whether it could increase the brain pericyte number or pericyte coverage of brain vessels on the larvae when the EC-specific Notch was inhibited (***Figure 5-figure supplementary 1C***). If blood flow works through EC-intrinsic Notch signaling, the number of brain pericytes or pericyte coverage of brain vessels should be unchanged in the above larvae. We found that neither up-regulation of blood flow nor increasing Piezo1 activity could increase the number of pericytes or pericyte coverage of vessels in the brain of *Tg(flk1:H2B-mNeoGreen-dnMAML)* larvae (number: NB = 24.2±1.7 versus DMSO = 24.4±2.3, p = 0.94, Yoda1 = 24.4±1.8 versus DMSO = 24.4±2.3, p = 0.98; coverage: NB = 0.35±0.07 versus DMSO = 0.31±0.10, p = 0.49, Yoda1 = 0.34±0.08 versus DMSO = 0.31±0.10, p = 0.43) (***Figure 5E*** and ***Figure 5-figure supplementary 1D***), demonstrating that EC–specific Notch inhibition could suppress the effects of NB or Yoda1 on the population expansion of brain pericytes. These results provide evidence that EC-intrinsic Notch signaling mediates the regulation of blood flow in brain pericyte proliferation.

## Discussion

Here we uncover a regulatory pathway that blood flow promotes brain pericyte proliferation through Piezo1-dependent up-regulation of Notch signaling in ECs, highlighting a role of hemodynamics for the regulation of perivascular cell development.

Besides the role in NVC, pericytes can also reinforce the rigidity of ECs and help the endothelial wall to withstand the blood flow-generating mechanical stress (***Armulik et al., 2011; Winkler et al., 2011***). Whereas our results here show that blood flow can also modulate brain pericyte proliferation, suggesting a mutual regulation between blood flow and brain pericytes for maintaining both of the blood supply and cerebrovascular integrity by a feedback pathway. Moreover, the high flow and low hydrodynamic resistance make the brain vasculature subjected to high pressure pulsatility ***(O’Rourke and Safar, 2005***), which can result in significant mechanical forces and consequent high Piezo1 activity in ECs (***Coste et al., 2010; Nourse and Pathak, 2017***). Our findings, combined with previous reports, suggest that the higher pericyte coverage of brain vasculature may be due to the unique hemodynamics-induced Piezo1 activity-dependent higher frequency of pericyte proliferation in the brain, compared to vasculature in other organs. Additionally, any physiological stimulus that affects hemodynamics, such as neural activity, could potentially regulate brain pericyte proliferation during development (***Andreone et al., 2015; Attwell et al., 2010***). Thus, our study underscores the essential need to consider the coordinated influences of both mechanical forces and chemical signals to gain a comprehensive understanding of brain pericyte development.

Piezo1 activation triggers Ca2+ signals in capillary ECs, which has profound effects on blood vessels (***Harraz et al., 2022***). Our *in vivo* results show that Notch signaling in brain ECs could be activated by the Piezo1 agonist Yoda1. Whereas how Notch signaling in ECs is activated by increased Piezo1 activity need to be explored in the future studies. Before localized on cell membrane, Notch receptor undergoes proteolysis by the Ca^2+^-dependent proteases inside the endoplasmic reticulum (ER) and Golgi apparatus before maturation, and disruption of Ca^2+^ homeostasis in the ER/Golgi impaired Notch receptor trafficking in cells (***Periz and Fortini, 1999***). Meanwhile, the activation of Notch requires the Ca^2+^-dependent ADAM metalloprotease-mediated cleavage ***(Mamaeva et al., 2009***). Loss of ADAM10 function was found to impair the cleavage of Notch1 in ECs and inhibit its intracellular Notch signaling transduction (***Caolo et al., 2020)***. Additionally, Ca^2+^ serves as a key second massager in multiple signaling pathways, some of which interact with Notch signaling. For example, VEGF functions upstream of Notch during vascular development (***Lawson et al., 2002***), and intercellular transduction of VEGF signaling is dependent on Ca^2+^ signaling by Ca^2+^/Calmodulin-dependent protein kinase II (***Banumathi et al., 2011***). In all, based on these findings, the increased Ca^2+^ could enhance Notch signaling through the activation of Notch receptor and/or the transduction of VEGF signaling upstream of Notch.

Although the Notch3 signaling was previously known to cell-autonomously regulate pericyte development (***Wang et al., 2014***), our findings indicate that the EC-intrinsic Notch signaling also promotes brain pericyte proliferation and mediates the effect of blood flow for brain pericyte development. However, it still unknown how the blood flow-evoked Notch signaling in ECs is transmitted to pericytes. Previous *in vitro* studies suggest that increasing Notch signaling in ECs led to increased expression of Notch ligands, such as Jag-1 (***Caolo et al., 2020; Chen et al., 2017***). And Jag-1 in turn could activate Notch3 receptor in pericytes to promote proliferation (***Liu et al., 2009; Volz et al., 2015; Wang et al., 2014***). These results suggest that blood flow-triggered EC-to-pericyte Notch signaling cascade may drive the proliferation of brain pericytes during cerebrovascular development (***Figure 5F***). Future work could focus on dissecting the detailed mechanism by which EC interact with pericyte, such as identifying the Notch ligands that can be up-regulated by EC-intrinsic Notch signaling for the activation of Notch3 signaling in pericytes during cerebrovascular development.

## Methods

### Resource lists

Drugs, protein reagents, kits and instruments used in this study are listed in ***Supplementary table 1***. The sequences of the primers and sgRNAs used in this study are listed in ***Supplementary table 2***.

### Zebrafish husbandry

The adult fish were raised in the automatic fish housing system (ESEN, Beijing, China) at the temperature of 28°C, with a 14 hour light/1 0 hour dark cycle. The embryos were kept in 10% Hank’s solution (140 mM NaCl, 5.4 mM KCl, 4.2 mM NaHCO_3_, 1.3 mM CaCl_2_, 1.0 mM MgSO_4_, 0.44 mM KH_2_PO_4_, and 0.25 mM Na_2_HPO_4_, with the pH maintained at 7.2) in an incubator under the same temperature and light: dark cycle as the adult. Methylene blue was applied in the Hank’s solution before 2.0 dpf. The handling of zebrafish used in the study was approved by the Center of Excellence in Brain Science and Intelligence Technology, Chinese Academy of Sciences (CEBSIT, CAS).The developmental stages of the zebrafish larvae used in this study ranged from 1.5 dpf to 7.5 dpf.

### Zebrafish strains

The following lines were used: *Ki(pdgfrb:GAL4-VP16)ion33d* abbreviated as *Ki(pdgfrb:GAL4-VP16); Tg(4xnrUAS:GFP)c369* abbreviated as *Tg(4×nrUAS:GFP)* (***Akitake et al., 2011***)*; Tg(fli1ep:dsredex)um13* abbreviated as *Tg(fli1a:DsRedEx*) (***Lorent et al., 2010***)*; Tg(UAS-E1b:Kaede)s1999t* abbreviated as *Tg(UAS:Kaede)* (***He et al., 2012***)*; Tg(UAS:mCherry)zf408* abbreviated as Tg(UAS:mCherry) **(Covassin et al., 2009**)*; Tg(Xla.Eef1a1:mAG-gmnn.100)rw0410h* abbreviated as *Tg(EF1a1:mAG-gmnn.100)* (***Sugiyama et al., 2009***)*; piezo1^ion89d^*abbreviated as *piezo1^-/-^* (***Liu et al., 2020***); *Tg(kdrl:Cre)s898* abbreviated as *Tg(flk1:Cre)* (***Bertrand et al., 2010****); Ki(piezo1-loxp,myl7:DsRed)ion204d* abbreviated as *Ki(piezo1-loxp); Tg(EPV.TP1-Mmu.Hbb:d2GFP)mw43* abbreviated as *Tg(Tp1:d2GFP)* (***Han et al., 2016***)*; Tg(kdrl:tetoff-H2B-mNeoGreen-dnMAML)ion253d* abbreviated as *Tg(flk1:tetoff-H2B-mNeoGreen-dnMAML);* and *Tg(kdrl:tetoff-H2B-mScarlet-NICD)ion252d* abbreviated as *Tg(flk1:tetoff-H2B-mScarlet-NICD);*

### Genomic DNA purification and genotyping

To obtain the genomic DNA templates for PCR, the frozen larvae or the tail samples of adult fish were subjected to cell lysis solution (10 mM Tris-HCl, 10 mM EDTA, 200 mM NaCl, 0.05% SDS) for 24 hours. The lysate was then precipitated with 70% ethanol, then draught for 10 min in the 60 ℃ metal bath, and then resuspended in distilled water. The concentration of the purified DNA was determined to 10 - 100 ng/uL with ddH_2_O. PCR was carried out with the 2x EsTaq Master Mix kit (CWBIO) and corresponding primers, the results of genotyping were displayed by electrophoresis or sequencing.

### Production of zCas9 mRNA, Tol2 mRNA and single guide RNA (sgRNA)

The XbaI linearized zebrafish codon-optimized Cas9 (zCas9) expression plasmid pGH-T7-zCas9 was used as a template for in vitro synthesis and purification of Cas9 mRNA, with the mMACHINE T7 Ultra kit (Ambion). The Tol2 mRNA was generated using the same protocol, with the plasmid pGH-T7-Tol2 as a template. The sgRNAs were designed using the previously reported criteria (***Chang et al., 2013***), aimed at selecting specific targets and minimizing off-target effects. A pair of oligonucleotides containing the sgRNA targeting sequence were annealed and cloned to the downstream of the T7 promoter in the PT7-sgRNA vector. The sgRNA was synthesized using the MAXIscript T7 Kit (Ambion) and purified by using the mirVana™ miRNA Isolation Kit (Ambion).

### Generation of knockin lines

The *Ki(pdgfrb:GAL4-VP16)* line used in this work was generated by CRISPR/Cas9 technology as previous report (***Li et al., 2015***). The optimal sgRNA target was selected through a mutation efficiency test for the sgRNA candidates. The donor plasmid was constructed by replacing the left and right arms of *th-P2A-Gal4-VP16* donor with corresponding arms of *pdgfrb* (***Li et al., 2015***). A drop of solution containing 800 pg Cas9 mRNAs, 100 pg sgRNAs and 15 pg donor plasmids was injected into the one-cell-stage embryos of *Tg(4×nrUAS:GFP)* line, the injected larvae that displayed green fluorescence were raised as F0 candidates. The F0 founder was obtained through outcrossing, and the fluorescence-positive F1 were raised for establishing the stable line. The correct insertion of the donor plasmid into the genome was confirmed using primers pdgfrb seq F1/R1 and pdgfrb seq F2/R2.

The *Ki(piezo1-loxp,myl7:DsRed)* line was generated using a similar protocol as the *Ki(pdgfrb:GAL4-VP16)* line. Specially, two efficient sgRNA targets were selected in the exon 24 of *piezo1*. The donor plasmid was constructed as figure S4, and consisted of the 1280 bp left arm, the flox exon 24 of *piezo1*, the 920 bp right arm and a co-marker *myl7:DsRed* to label the heart as red fluorescence. After the one-cell-stage injection, the larvae that displayed *myl7:DsRed* labeled in their heart were raised as F0 candidates. The founder was screened through observation of heart fluorescence and PCR for the offspring.

### Generation of transgenic lines

*Tg(kdrl:tetoff-H2B-mScarlet-NICD) and Tg(kdrl:tetoff-H2B-mNeoGreen-dnMAML)* line were generated by using Tol2 mediated transgenesis (***Kawakami et al., 2016***). Firstly, to generate the plasmids, the cDNAs of NICD and dnMAML were amplified from the zebrafish cDNA library with the corresponding primers, and the DNA fragments of tetoff, H2B-mScarlet, H2B-mNeoGreen were amplified from relevant template plasmids, then DNA fragments were cloned into the backbones containing Tol2-flk, and finally the plasmids Tol2-flk1-tetoff-H2B-mScarlet-NICD, Tol2-flk1-tetoff-H2B-mNeoGreen-dnMAML were identified by colony PCR and DNA sequencing. To generate the transgenic lines, a drop of solution containing 25 pg each plasmid and 25 pg Tol2 mRNA was injected to the one-cell embryos, and the larvae with fluorescence expression were kept as F0 candidates, and the founder was screened by the fluorescence pattern of the offspring.

### *In vivo* imaging and cell fate tracking of zebrafish larvae

*In vivo* fluorescent images of the zebrafish embryos were captured at room temperature using an Olympus FV1000 or FV3000 confocal microscope. Live embryos were mounted in 1.5% low-melting agarose (Invitrogen) containing 0.2% MS-222 in 35mm glass bottom petri dishes (MatTek). The images were captured with a z-step of 3 to 5 micrometers, and had a resolution of either 1024 x 1024 or 512 x 512 pixels. The ImageJ software was utilized to analyze these images.

Photoconversion was carried out with the FV1000 confocal fluorescence microscopy at 2.5 dpf with the larvae mounted in 1.5% low-melting agarose (Invitrogen). The region containing the single pericyte of interest was selected with the zoom function, and exposed to 405 nm laser for 10-20 seconds. Then the whole brain *in vivo* imaging was quickly carried out to record the start state of the photoconverted brain pericyte. The larvae were continuously raised in dark condition, and the second whole brain imaging was performed at 4.5 dpf to track the fate of the photoconverted pericytes.

### Drug application

The BDM stock was stored at 4 M in DMSO, and 4 uM of it was used to decrease blood flow. The NB stock was stored at 50 mM in ddH2O, and 250 uM of it was used to increase blood flow. The stock of Yoda1 was stored at 10 mM in DMSO, and 1 uM of it was used to activate Piezo1. The stock of GsMTx4 was stored at 0.5 mM in ddH2O, and 1 uM of it was used to inhibit Piezo1 activity. The stock of DAPT was stored at 50 mM in DMSO, and 50 uM of it was used to inhibit overall Notch signaling. All the drugs in this study were applied during 3.5 - 4.5 dpf for counting the brain pericyte number and pericyte coverage of brain vessels, and used during 3.0 - 3.5 dpf for time-lapse imaging to count the proliferation frequency of brain pericytes and event number of brain ingress of the pericerebral pericytes.

### Blood flow assessment

The velocity of blood flow was measured by recording the heart rate of each larva. To do this, the larva was placed in a drop of 10% Hank’s solution, and its heart beating number was recorded over a period of 30 seconds. The final heat rate value was defined as the number of heart beats per minute.

### Quantification of pericyte coverage of brain vessels and brain pericyte proliferation frequency

The cell counter function of ImageJ was used to manually count the number of pericytes and vessel segments in the brain of the larvae. The pericyte coverage of brain vessels was then calculated as the average number of pericytes per vessel segment (***Xu et al., 2017***). The number of starting brain pericytes and the increased brain pericytes were counted with the time-lapse images collected during 3.0 - 3.5 dpf, and the mean proliferation frequency of brain pericytes was calculated as the total number of brain pericyte proliferation events occurred during the 12 hours divided by the number of brain pericytes at 3.0 dpf. The analysis was performed blindly without prior knowledge of the treatment or genotype of the larvae.

### Statistics

Zebrafish larvae were assigned to experimental and control groups randomly, without investigator blinding. The significance of the difference between two groups was calculated using an unpaired two-tailed Student’s t-test, which was performed using the GraphPad Prism v9.0 software. The results were represented as mean±SEM in all figures, and the level of significance was indicated as follows: **** p < 0.0001, *** p < 0.001, ** indicates p < 0.01, * indicates p < 0.05, and n.s. indicates p > 0.05.

## Acknowledgments

We thank Dr. Qian Hu for helping confocal imaging and Dr. Jie He for providing the *Tg(UAS:Kaede)* and *Tg(EF1a1:mAG-gmnn.100)* lines. This work was supported by the National Key R&D Program of China (No. 2019YFA0801603 to J.L.), National Natural Science Foundation of China (NO. 31701265 to X.P.), and Shanghai Natural Science Foundation (NO.22ZR1469700 to J.L.).

## Author contributions

J.D., H.Z. and J.L. conceptualized and supervised the study; H.Z. and J.L. designed and performed the major experiments; J.L., H,Z., X.P. and J.B. generated and confirmed the *Ki(pdgfrb:GAL4-VP16)* line; H,Z. and J.C explored the proper application concentrations of NB, BDM and Yoda1; H,Z. and J.L. wrote the manuscript with inputs from J.D.

## Declaration of interests

The authors declare no competing interests.

---

**Figure 1-figure supplementary 1.**
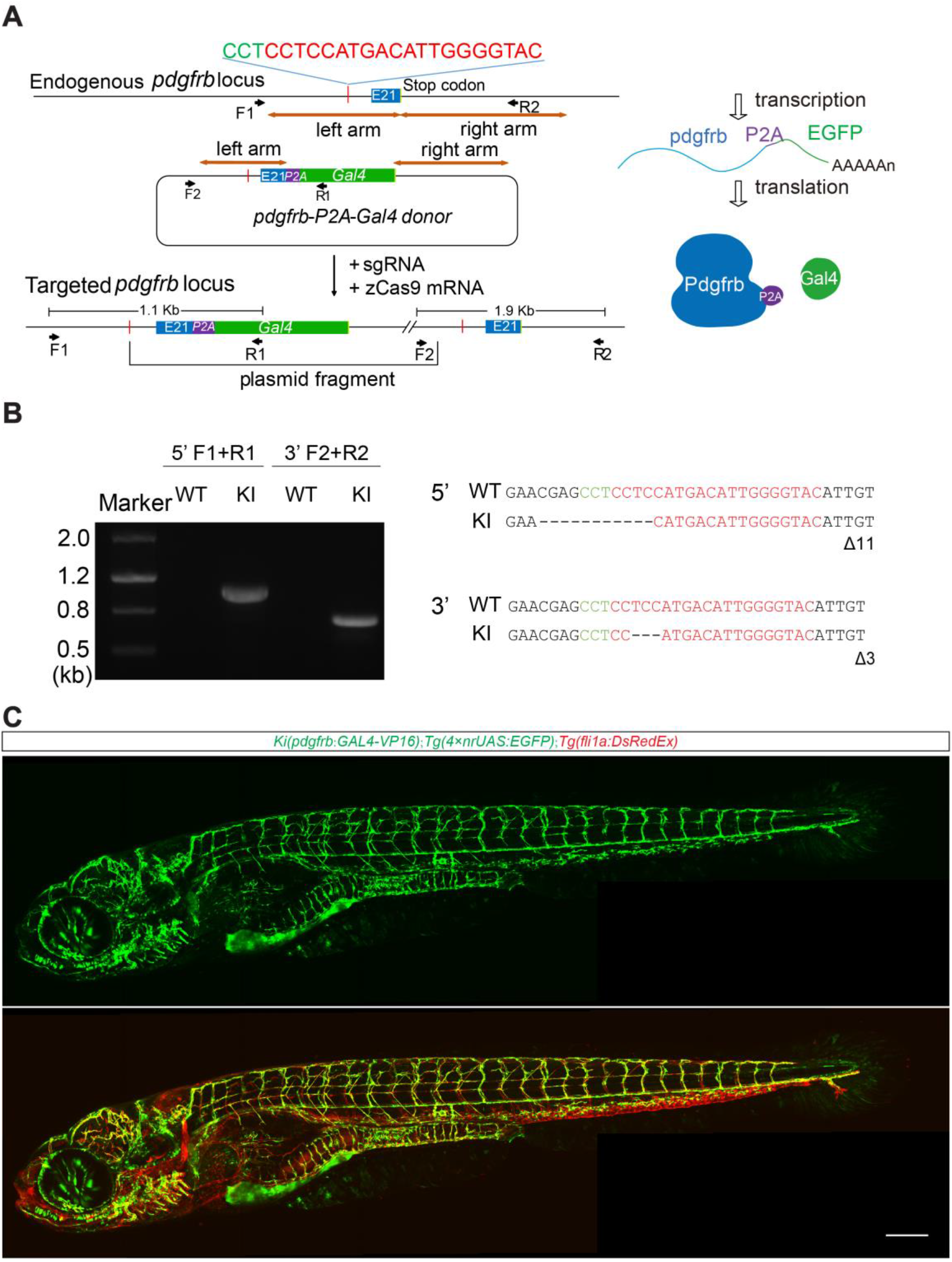
Generation of pericyte reporter line. (A) Schematic representation of the generation of the *Ki(pdgfrb:GAL4-VP16)* line using CRISPR/Cas9-mediated knock-in technology. (B) PCR and sequencing results for the 5’ and 3’ junction of donor plasmid integration into *pdgfrb* locus. (C) Example showing the ingress of pericerebral pericytes into the brain. Images are shown from a lateral view. Scale bar, 200 μm.

**Figure 1-figure supplementary 2.**
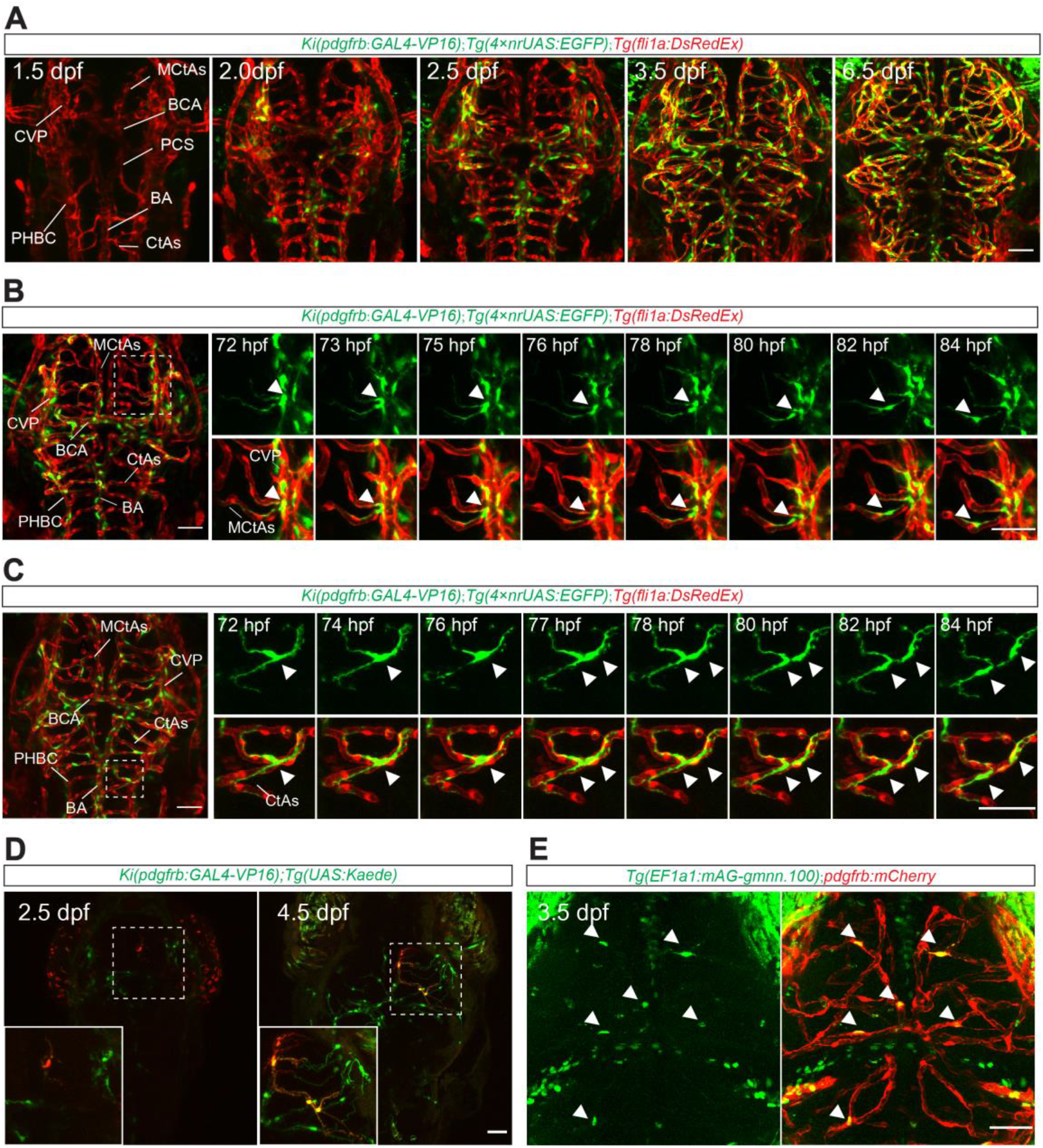
Developmental dynamics of brain pericytes during embryogenesis. (A) Images of the pericytes and blood vessels in the brain images of a *Ki(pdgfrb:GAL4-VP16);Tg(4×nrUAS:GFP);Tg(fli1a:DsRedEx)* larva from 1.5 dpf to 6.5 dpf. (B) Example showing the ingress of pericytes from the pericerebral vessels to the brain. (C) Example showing the proliferation of pericytes in the brain. (D) Example showing cell fate tracking for brain pericytes. A pericyte in the midbrain of a *Ki(pdgfrb:GAL4-VP16);Tg(UAS:Kaede)* larva was marked with red fluorescence via photo-conversion at 2.5 dpf, and its daughter cells that expressing red fluorescence were observed at 4.5 dpf. (E) Representative brain image of the *Tg(EF1a1:mAG-gmnn.100);Ki(pdgfrb:GAL4-VP16);Tg(UAS:mCherry-MA)* larvae at 3.5 dpf. The brain pericytes expressing green fluorescence are marked with arrowheads. *Ki(pdgfrb:GAL4-VP16);Tg(UAS:mCherry-MA)* abbreviated to *pdgfrb:mCherry* here. Images are shown from a top view. Scale bar, 50 μm for (A-E).

**Figure 1-figure supplementary 3.**
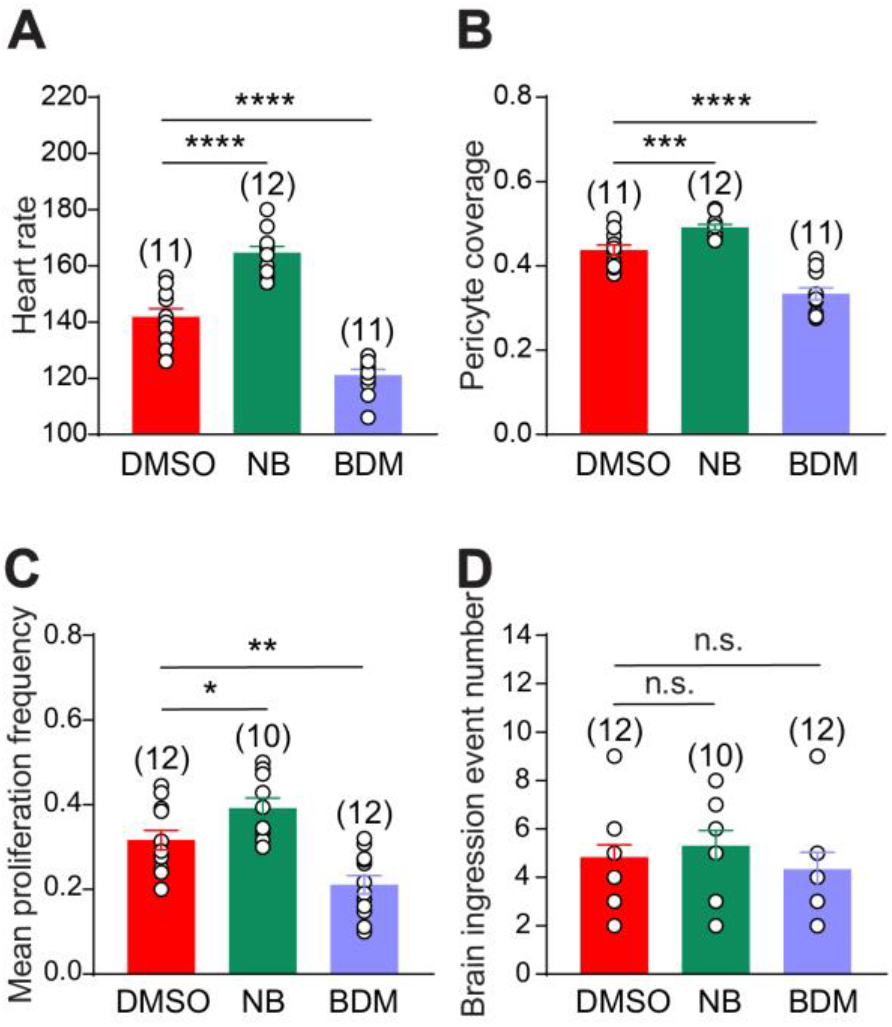
Blood flow promotes brain pericyte proliferation. (A) Summary of the effects of BDM and NB treatment on blood flow of 4.5 dpf larvae. The blood flow velocity is indicated by the heart rate. (B-D) Summary of the effects of BDM and NB treatment on pericyte coverage of brain vessels (B), brain pericyte proliferation (C), and brain ingression of pericerebral pericytes (D). Data are represented as mean ± SEM. N values are shown above the aligned plots. Images of 4.5 dpf larvae are used for counting the brain pericyte number and the pericyte coverage of brain vessels, time-lapse images during 3.0-3.5 dpf are used for counting the event numbers of brain pericyte proliferation and brain ingress of the pericerebral pericytes. Stars represent the results of unpaired two-tailed Student’s t-test between groups (*p < 0.05, **p < 0.01, ***p < 0.001, ****p < 0.0001).

**Figure 2-figure supplementary 1.**
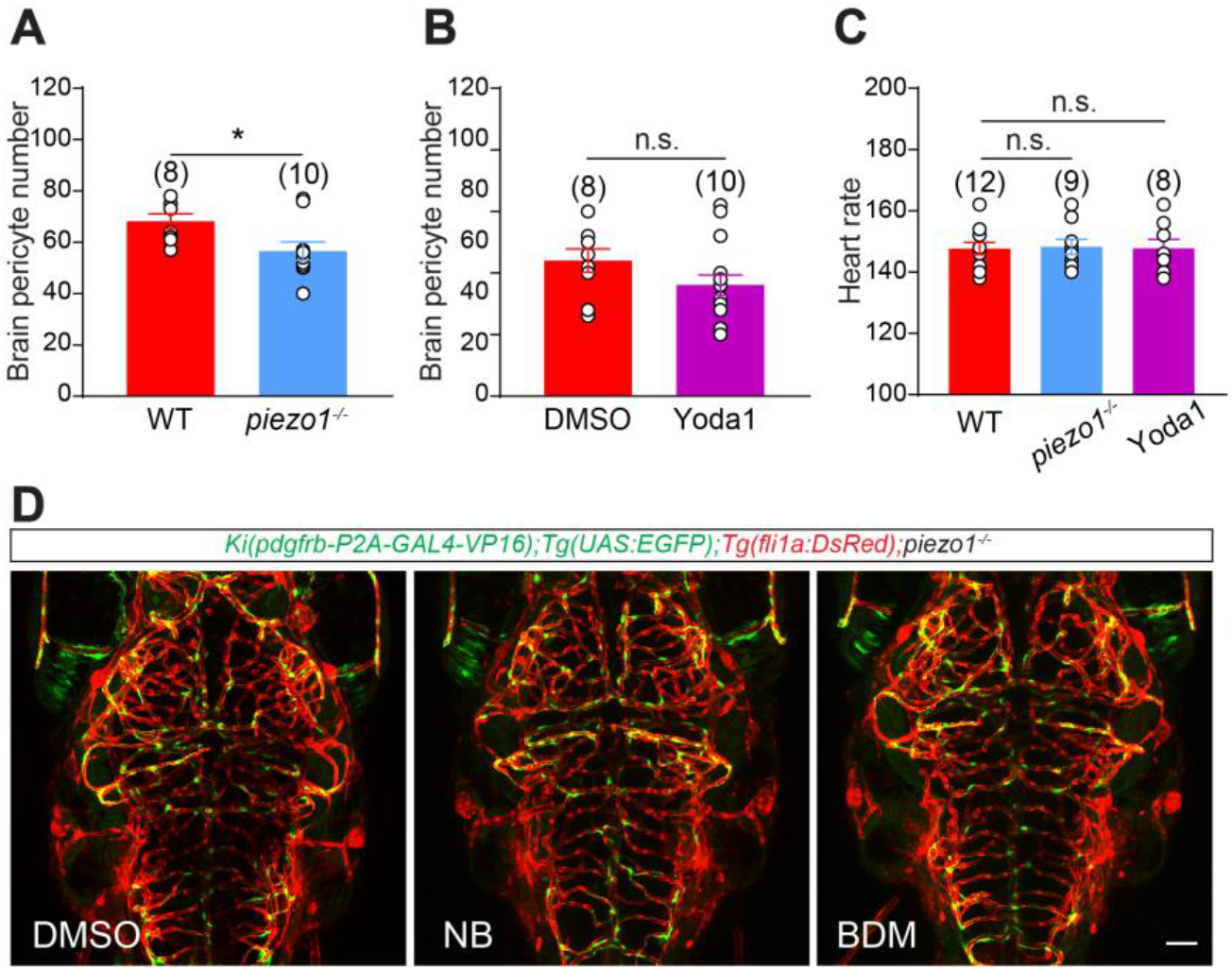
Piezo1 mediates the effect of blood flow on brain pericyte proliferation. (A) Summary of the effect of *piezo1* knockout on brain pericyte number. (B) Summary of the effect of Yoda1 treatment on brain pericyte number. (C) Graph showing the heart rates of the WT, *piezo1^-/-^* and the Yoda1 treated larvae at 4.5 dpf. (D) Representative images of the pericytes and blood vessels in the brain of DMSO, NB, and Yoda1 treated *piezo1^-/-^*larvae at 4.5 dpf. Images are shown from a top view. Scale bar, 50 μm for (D). Images of 4.5 dpf larvae are used for counting the brain pericyte number and the pericyte coverage of brain vessels. Data are represented as mean ± SEM. N values are shown above the aligned plots. Stars represent the results of unpaired two-tailed Student’s t-test between groups (*p < 0.05, **p < 0.01, ***p < 0.001, ****p < 0.0001).

**Figure 3-figure supplementary 1.**
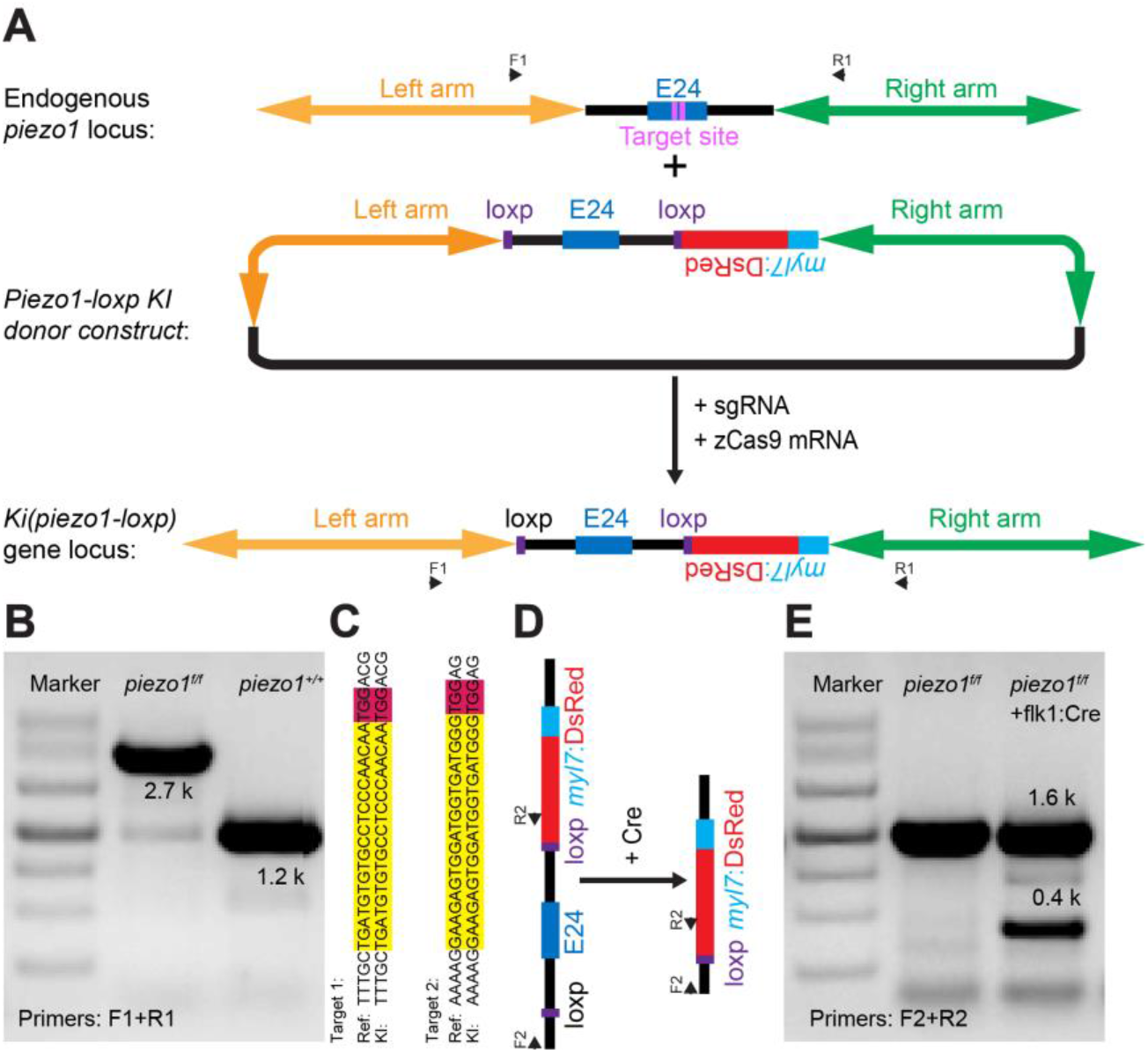
Generation of *Ki(piezo1-loxp)* line. (A) Schematic illustrating the CRISPR/Cas9-mediated knock-in for *Ki(piezo1-loxp)* line. The *myl7:DsRed* enables the stable line express red fluorescence DsRed in the heart as a co-marker. (B-E) PCR and sequencing for confirming the targeted integration of Loxp into the *piezo1* locus.

**Figure 4-figure supplementary 1.**
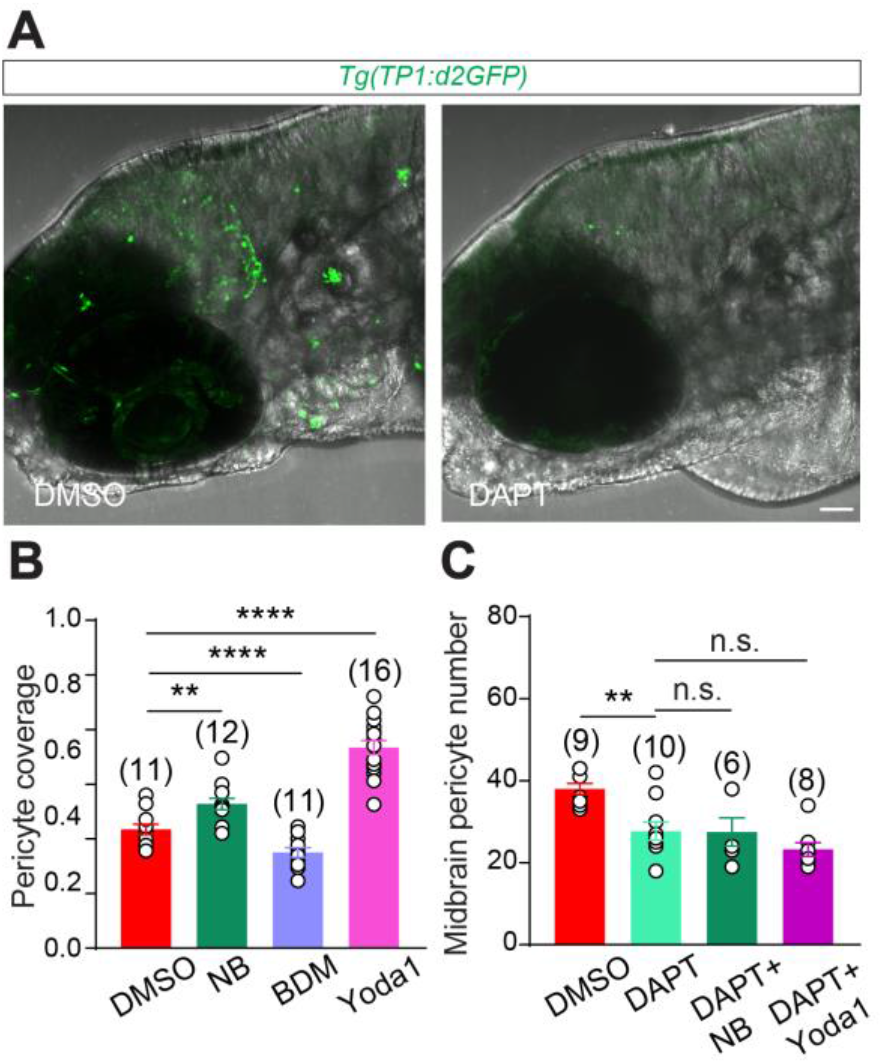
Notch signaling functions as the downstream of Piezo1 in regulating pericyte proliferation. (A) Representative images showing the fluorescent pattern in the brain of *Tg(Tp1:d2GFP)* larvae and the effect of DAPT treatment on it. DAPT was applied during 3.0 - 4.0 dpf, and the imaging was performed at 4.0 dpf. (B) Summary of the effects of BDM, NB and Yoda1 treatment on pericyte coverage of vessels in the midbrain. (C) Summary of the effect of Yoda1 treatment on the pericyte coverage of vessels in the midbrain. Images are shown from a top view. Scale bar, 50 μm for (A). Images of 4.5 dpf larvae are used for counting the pericyte coverage of vessels in the midbrain. Data are represented as mean ± SEM. The n values are shown above the aligned plots. Stars represent the results of unpaired two-tailed Student’s t-test between groups (*p < 0.05, **p < 0.01, ***p < 0.001, ****p < 0.0001).

**Figure 5-figure supplementary 1.**
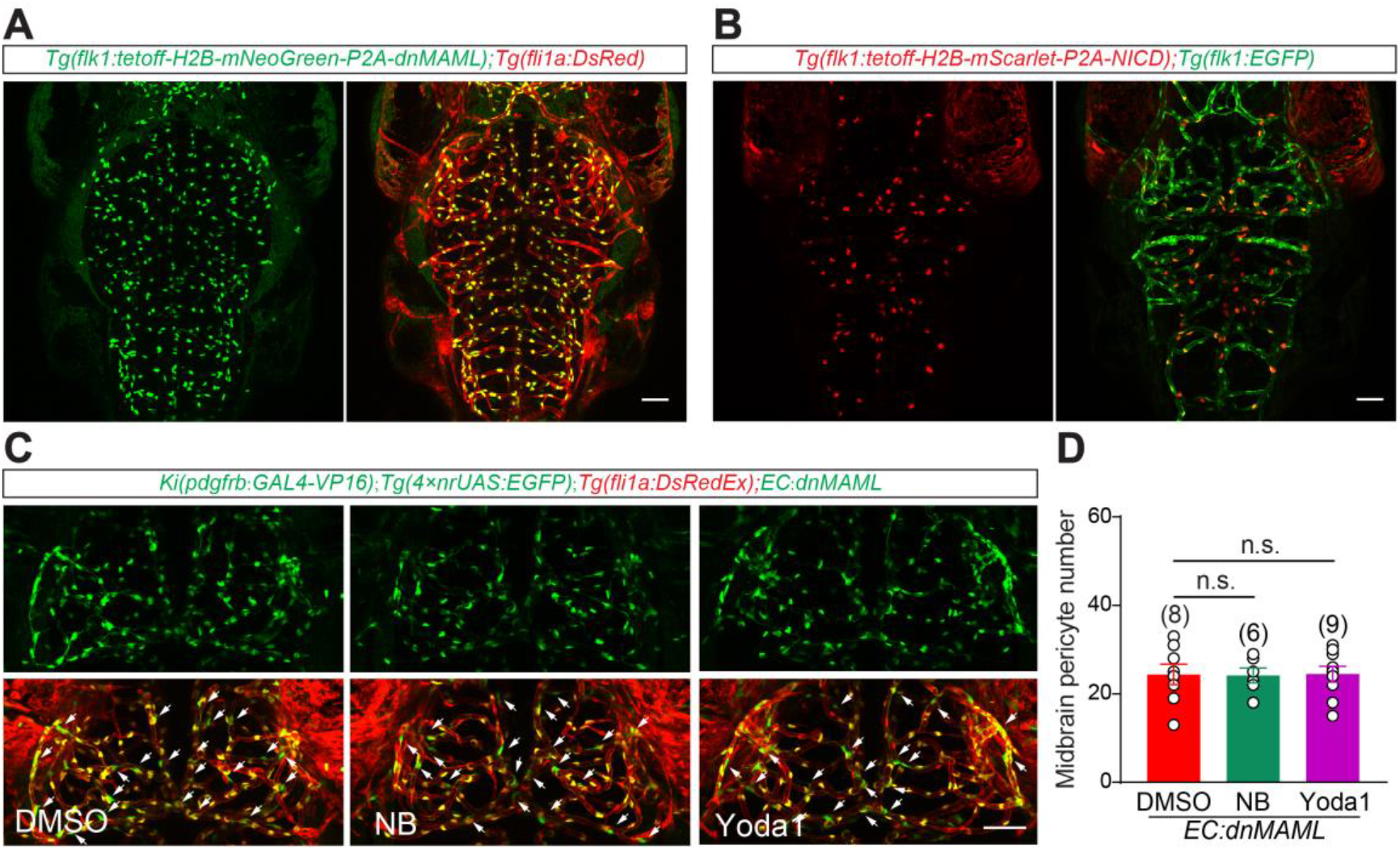
EC-specific Notch signaling mediates blood flow regulation of brain pericyte proliferation. (A) Representative images of the blood vessels in the brain of *Tg(fli1a:DsRedEx)*;*Tg(flk1:tetoff-H2B-mNeoGreen-P2A-dnMAML)* larvae at 4.5 dpf. (B) Representative images of the blood vessels in the brain of *Tg(flk1:EGFP);Tg(flk1: tetoff-H2B-mScarlet-P2A-NICD)* larvae at 4.5 dpf. (C) Representative images of the pericytes and blood vessels in the midbrain of DMSO, NB and Yoda1 treated larvae at 4.5 dpf. (D) Summary of the effects of DMSO, NB and Yoda1 treatment on the number of midbrain pericytes in the *Tg(flk1:tetoff-H2B-mNeoGreen-P2A-dnMAML)* larvae, which is abbreviated as *EC: dnMAML* here. Images are shown from a top view. Scale bar, 50 μm for (A-C). Images of 4.5 dpf larvae are used for counting the number of pericytes in the midbrain. Data are represented as mean ±SEM. The n values are shown above the aligned plots. Stars represent the results of unpaired two-tailed Student’s t-test between groups (*p < 0.05, **p < 0.01, ***p < 0.001, ****p < 0.0001).

## Notes

### Competing Interest Statement

The authors have declared no competing interest.

### Summary of Updates

we modified the author order shown on the website, no change of it in the PDF file

## Reference

Akitake, C.M., Macurak, M., Halpern, M.E., and Goll, M.G. (2011). Transgenerational analysis of transcriptional silencing in zebrafish. Developmental biology 352, 191–201.

Alarcon-Martinez, L., Villafranca-Baughman, D., Quintero, H., Kacerovsky, J.B., Dotigny, F., Murai, K.K., Prat, A., Drapeau, P., and Di Polo, A. (2020). Interpericyte tunnelling nanotubes regulate neurovascular coupling. Nature 585, 91–95.

Ando, K., Fukuhara, S., Izumi, N., Nakajima, H., Fukui, H., Kelsh, R.N., and Mochizuki, N. (2016). Clarification of mural cell coverage of vascular endothelial cells by live imaging of zebrafish. Development 143, 1328–1339.

Andreone, B.J., Lacoste, B., and Gu, C. (2015). Neuronal and vascular interactions. Annual review of neuroscience 38, 25–46.

Armulik, A., Genove, G., and Betsholtz, C. (2011). Pericytes: developmental, physiological, and pathological perspectives, problems, and promises. Developmental cell 21, 193–215.

Armulik, A., Genove, G., Mae, M., Nisancioglu, M.H., Wallgard, E., Niaudet, C., He, L., Norlin, J., Lindblom, P., Strittmatter, K., et al. (2010). Pericytes regulate the blood-brain barrier. Nature 468, 557–561.

Attwell, D., Buchan, A.M., Charpak, S., Lauritzen, M., Macvicar, B.A., and Newman, E.A. (2010). Glial and neuronal control of brain blood flow. Nature 468, 232–243.

Banumathi, E., O’Connor, A., Gurunathan, S., Simpson, D.A., McGeown, J.G., and Curtis, T.M. (2011). VEGF-induced retinal angiogenic signaling is critically dependent on Ca(2)(+) signaling by Ca(2)(+)/calmodulin-dependent protein kinase II. Investigative ophthalmology & visual science 52, 3103–3111.

Bertrand, J.Y., Chi, N.C., Santoso, B., Teng, S., Stainier, D.Y., and Traver, D. (2010). Haematopoietic stem cells derive directly from aortic endothelium during development. Nature 464, 108–111.

Botello-Smith, W.M., Jiang, W., Zhang, H., Ozkan, A.D., Lin, Y.C., Pham, C.N., Lacroix, J.J., and Luo, Y. (2019). A mechanism for the activation of the mechanosensitive Piezo1 channel by the small molecule Yoda1. Nature communications 10, 4503.

Caolo, V., Debant, M., Endesh, N., Futers, T.S., Lichtenstein, L., Bartoli, F., Parsonage, G., Jones, E.A., and Beech, D.J. (2020). Shear stress activates ADAM10 sheddase to regulate Notch1 via the Piezo1 force sensor in endothelial cells. eLife 9.

Chang, N., Sun, C., Gao, L., Zhu, D., Xu, X., Zhu, X., Xiong, J.W., and Xi, J.J. (2013). Genome editing with RNA-guided Cas9 nuclease in zebrafish embryos. Cell research 23, 465–472.

Chen, Q., Jiang, L., Li, C., Hu, D., Bu, J.W., Cai, D., and Du, J.L. (2012). Haemodynamics-driven developmental pruning of brain vasculature in zebrafish. PLoS biology 10, e1001374.

Chen, X., Gays, D., Milia, C., and Santoro, M.M. (2017). Cilia Control Vascular Mural Cell Recruitment in Vertebrates. Cell reports 18, 1033–1047.

Clark, B.S., Cui, S., Miesfeld, J.B., Klezovitch, O., Vasioukhin, V., and Link, B.A. (2012). Loss of Llgl1 in retinal neuroepithelia reveals links between apical domain size, Notch activity and neurogenesis. Development 139, 1599–1610.

Coste, B., Mathur, J., Schmidt, M., Earley, T.J., Ranade, S., Petrus, M.J., Dubin, A.E., and Patapoutian, A. (2010). Piezo1 and Piezo2 are essential components of distinct mechanically activated cation channels. Science 330, 55–60.

Covassin, L.D., Siekmann, A.F., Kacergis, M.C., Laver, E., Moore, J.C., Villefranc, J.A., Weinstein, B.M., and Lawson, N.D. (2009). A genetic screen for vascular mutants in zebrafish reveals dynamic roles for Vegf/Plcg1 signaling during artery development. Developmental biology 329, 212–226.

Daneman, R., Zhou, L., Kebede, A.A., and Barres, B.A. (2010). Pericytes are required for blood-brain barrier integrity during embryogenesis. Nature 468, 562–566.

Duchemin, A.L., Vignes, H., and Vermot, J. (2019). Mechanically activated piezo channels modulate outflow tract valve development through the Yap1 and Klf2-Notch signaling axis. eLife 8.

Galvez-Santisteban, M., Chen, D., Zhang, R., Serrano, R., Nguyen, C., Zhao, L., Nerb, L., Masutani, E.M., Vermot, J., Burns, C.G., et al. (2019). Hemodynamic-mediated endocardial signaling controls in vivo myocardial reprogramming. eLife 8.

Geling, A., Steiner, H., Willem, M., Bally-Cuif, L., and Haass, C. (2002). A gamma-secretase inhibitor blocks Notch signaling in vivo and causes a severe neurogenic phenotype in zebrafish. EMBO reports 3, 688–694.

Geudens, I., Coxam, B., Alt, S., Gebala, V., Vion, A.C., Meier, K., Rosa, A., and Gerhardt, H. (2019). Artery-vein specification in the zebrafish trunk is pre-patterned by heterogeneous Notch activity and balanced by flow-mediated fine-tuning. Development 146.

Hahn, C., and Schwartz, M.A. (2009). Mechanotransduction in vascular physiology and atherogenesis. Nature reviews Molecular cell biology 10, 53–62.

Han, P., Bloomekatz, J., Ren, J., Zhang, R., Grinstein, J.D., Zhao, L., Burns, C.G., Burns, C.E., Anderson, R.M., and Chi, N.C. (2016). Coordinating cardiomyocyte interactions to direct ventricular chamber morphogenesis. Nature 534, 700–704.

Harraz, O.F., Klug, N.R., Senatore, A.J., Hill-Eubanks, D.C., and Nelson, M.T. (2022). Piezo1 Is a Mechanosensor Channel in Central Nervous System Capillaries. Circulation research 130, 1531–1546.

He, J., Zhang, G., Almeida, A.D., Cayouette, M., Simons, B.D., and Harris, W.A. (2012). How variable clones build an invariant retina. Neuron 75, 786–798.

Kawakami, K., Asakawa, K., Muto, A., and Wada, H. (2016). Tol2-mediated transgenesis, gene trapping, enhancer trapping, and Gal4-UAS system. Methods in cell biology 135, 19–37.

Kisler, K., Nelson, A.R., Rege, S.V., Ramanathan, A., Wang, Y., Ahuja, A., Lazic, D., Tsai, P.S., Zhao, Z., Zhou, Y., et al. (2017). Pericyte degeneration leads to neurovascular uncoupling and limits oxygen supply to brain. Nature neuroscience.

Lasch, M., Kleinert, E.C., Meister, S., Kumaraswami, K., Buchheim, J.I., Grantzow, T., Lautz, T., Salpisti, S., Fischer, S., Troidl, K., et al. (2019). Extracellular RNA released due to shear stress controls natural bypass growth by mediating mechanotransduction in mice. Blood 134, 1469–1479.

Lawson, N.D., Vogel, A.M., and Weinstein, B.M. (2002). sonic hedgehog and vascular endothelial growth factor act upstream of the Notch pathway during arterial endothelial differentiation. Developmental cell 3, 127–136.

le Noble, F., Klein, C., Tintu, A., Pries, A., and Buschmann, I. (2008). Neural guidance molecules, tip cells, and mechanical factors in vascular development. Cardiovascular research 78, 232–241.

le Noble, F., Moyon, D., Pardanaud, L., Yuan, L., Djonov, V., Matthijsen, R., Breant, C., Fleury, V., and Eichmann, A. (2004). Flow regulates arterial-venous differentiation in the chick embryo yolk sac. Development 131, 361–375.

Li, J., Hou, B., Tumova, S., Muraki, K., Bruns, A., Ludlow, M.J., Sedo, A., Hyman, A.J., McKeown, L., Young, R.S., et al. (2014). Piezo1 integration of vascular architecture with physiological force. Nature 515, 279–282.

Li, J., Zhang, B.B., Ren, Y.G., Gu, S.Y., Xiang, Y.H., Huang, C., and Du, J.L. (2015). Intron targeting-mediated and endogenous gene integrity-maintaining knockin in zebrafish using the CRISPR/Cas9 system. Cell research.

Lindahl, P., Johansson, B.R., Leveen, P., and Betsholtz, C. (1997). Pericyte loss and microaneurysm formation in PDGF-B-deficient mice. Science 277, 242–245.

Liu, H., Kennard, S., and Lilly, B. (2009). NOTCH3 expression is induced in mural cells through an autoregulatory loop that requires endothelial-expressed JAGGED1. Circulation research 104, 466–475.

Liu, H., Zhang, W., Kennard, S., Caldwell, R.B., and Lilly, B. (2010). Notch3 is critical for proper angiogenesis and mural cell investment. Circulation research 107, 860–870.

Liu, T.T., Du, X.F., Zhang, B.B., Zi, H.X., Yan, Y., Yin, J.A., Hou, H., Gu, S.Y., Chen, Q., and Du, J.L. (2020). Piezo1-Mediated Ca(2+) Activities Regulate Brain Vascular Pathfinding during Development. Neuron 108, 180–192 e185.

Lorent, K., Moore, J.C., Siekmann, A.F., Lawson, N., and Pack, M. (2010). Reiterative use of the notch signal during zebrafish intrahepatic biliary development. Developmental dynamics: an official publication of the American Association of Anatomists 239, 855–864.

Mamaeva, O.A., Kim, J., Feng, G., and McDonald, J.M. (2009). Calcium/calmodulin-dependent kinase II regulates notch-1 signaling in prostate cancer cells. Journal of cellular biochemistry 106, 25–32.

Mishra, A., Reynolds, J.P., Chen, Y., Gourine, A.V., Rusakov, D.A., and Attwell, D. (2016). Astrocytes mediate neurovascular signaling to capillary pericytes but not to arterioles. Nature neuroscience 19, 1619–1627.

Nakajima, H., Yamamoto, K., Agarwala, S., Terai, K., Fukui, H., Fukuhara, S., Ando, K., Miyazaki, T., Yokota, Y., Schmelzer, E., et al. (2017). Flow-Dependent Endothelial YAP Regulation Contributes to Vessel Maintenance. Developmental cell 40, 523–536 e526.

Nourse, J.L., and Pathak, M.M. (2017). How cells channel their stress: Interplay between Piezo1 and the cytoskeleton. Seminars in cell & developmental biology 71, 3–12.

O’Rourke, M.F., and Safar, M.E. (2005). Relationship between aortic stiffening and microvascular disease in brain and kidney: cause and logic of therapy. Hypertension 46, 200–204.

Obi, S., Yamamoto, K., Shimizu, N., Kumagaya, S., Masumura, T., Sokabe, T., Asahara, T., and Ando, J. (2009). Fluid shear stress induces arterial differentiation of endothelial progenitor cells. Journal of applied physiology 106, 203–211.

Pan, X., Wan, H., Chia, W., Tong, Y., and Gong, Z. (2005). Demonstration of site-directed recombination in transgenic zebrafish using the Cre/loxP system. Transgenic research 14, 217–223.

Periz, G., and Fortini, M.E. (1999). Ca(2+)-ATPase function is required for intracellular trafficking of the Notch receptor in Drosophila. The EMBO journal 18, 5983–5993.

Ranade, S.S., Qiu, Z., Woo, S.H., Hur, S.S., Murthy, S.E., Cahalan, S.M., Xu, J., Mathur, J., Bandell, M., Coste, B., et al. (2014). Piezo1, a mechanically activated ion channel, is required for vascular development in mice. Proceedings of the National Academy of Sciences of the United States of America 111, 10347–10352.

Ross Nortley, D.A. (2019). Amyloid β oligomers constrict human capillaries in Alzheimer’s disease via signaling to pericytes. Science

Sagare, A.P., Bell, R.D., Zhao, Z., Ma, Q., Winkler, E.A., Ramanathan, A., and Zlokovic, B.V. (2013). Pericyte loss influences Alzheimer-like neurodegeneration in mice. Nature communications 4, 2932.

Samsa, L.A., Givens, C., Tzima, E., Stainier, D.Y., Qian, L., and Liu, J. (2015). Cardiac contraction activates endocardial Notch signaling to modulate chamber maturation in zebrafish. Development 142, 4080–4091.

Schuster, K., and Ghysen, A. (2013). Labeling defined cells or subsets of cells in zebrafish by Kaede photoconversion. Cold Spring Harbor protocols 2013.

Sugiyama, M., Sakaue-Sawano, A., Iimura, T., Fukami, K., Kitaguchi, T., Kawakami, K., Okamoto, H., Higashijima, S., and Miyawaki, A. (2009). Illuminating cell-cycle progression in the developing zebrafish embryo. Proceedings of the National Academy of Sciences of the United States of America 106, 20812–20817.

Vanlandewijck, M., He, L., Mae, M.A., Andrae, J., Ando, K., Del Gaudio, F., Nahar, K., Lebouvier, T., Lavina, B., Gouveia, L., et al. (2018). A molecular atlas of cell types and zonation in the brain vasculature. Nature 554, 475–480.

Volz, K.S., Jacobs, A.H., Chen, H.I., Poduri, A., McKay, A.S., Riordan, D.P., Kofler, N., Kitajewski, J., Weissman, I., and Red-Horse, K. (2015). Pericytes are progenitors for coronary artery smooth muscle. eLife 4.

Wang, Y., Pan, L., Moens, C.B., and Appel, B. (2014). Notch3 establishes brain vascular integrity by regulating pericyte number. Development 141, 307–317.

Winkler, E.A., Bell, R.D., and Zlokovic, B.V. (2011). Central nervous system pericytes in health and disease. Nature neuroscience 14, 1398–1405.

Xu, B., Zhang, Y., Du, X.F., Li, J., Zi, H.X., Bu, J.W., Yan, Y., Han, H., and Du, J.L. (2017). Neurons secrete miR-132-containing exosomes to regulate brain vascular integrity. Cell research 27, 882–897.

Yang, A.C., Vest, R.T., Kern, F., Lee, D.P., Agam, M., Maat, C.A., Losada, P.M., Chen, M.B., Schaum, N., Khoury, N., et al. (2022). A human brain vascular atlas reveals diverse mediators of Alzheimer’s risk. Nature.

Yao, D., Zhang, R., Xie, M., Ding, F., Wang, M., and Wang, W. (2023). Updated Understanding of the Glial-Vascular Unit in Central Nervous System Disorders. Neuroscience bulletin 39, 503–518.

Zhao, Z., Nelson, A.R., Betsholtz, C., and Zlokovic, B.V. (2015). Establishment and Dysfunction of the Blood-Brain Barrier. Cell 163, 1064–1078.

